# Interaction of 14-3-3I and CDPK1 mediates the growth of human malaria parasite

**DOI:** 10.1101/2020.01.14.906479

**Authors:** Ravi Jain, Pinki Dey, Sakshi Gupta, Soumya Pati, Arnab Bhattacherjee, Manoj Munde, Shailja Singh

## Abstract

Scaffold proteins play pivotal role as modulators of cellular processes by operating as multipurpose conformation clamps. 14-3-3 proteins are gold-standard scaffold modules that recognize phosphoSer/Thr (*p*S/*p*T) containing conserved motifs of target proteins and confer conformational changes leading to modulation of their functional parameters. Modulation in functional activity of kinases has been attributed to their interaction with 14-3-3 proteins. Herein, we have characterized *Plasmodium falciparum* 14-3-3 and its interaction with key kinase of the parasite, Calcium-Dependent Protein Kinase 1 (CDPK1) by performing various analytical biochemistry and biophysical assays. Towards this, we annotated PF3D7_0818200 as 14-3-3 isoform I through extensive phylogenetic and comparative sequence analysis. Molecular dynamics simulation studies indicated that phosphoSer^64^ present in CDPK1 polypeptide sequence (^61^KLG*p*S^64^) behaves as canonical Mode I-type (RXX*p*S/*p*T) consensus 14-3-3 binding motif, mediating the interaction. The protein-protein interaction was validated *in vitro* with ELISA and SPR, which confirmed that CDPK1 interacts with 14-3-3I in a phosphorylation dependent manner, with binding affinity constant of 670 ± 3.6 nM. The interaction of 14-3-3I with CDPK1 was validated with well characterized optimal 14-3-3 recognition motifs: ARSH*p*SYPA and RLYH*p*SLPA as CDPK1 mimetics, by simulation studies and ITC. Further, interaction antagonizing peptidomimetics showed growth inhibitory impact on the parasite indicating crucial physiological role of 14-3-3/CDPK1 interaction. Overall, this study characterizes 14-3-3I as a scaffold protein in the malaria parasite and unveils CDPK1 as its previously unidentified target. This sets a precedent for the rational design of 14-3-3 based PPI inhibitors by utilizing 14-3-3 recognition motif peptides, as a potential antimalarial strategy.

---

Cellular signal transduction pathways often involve Post-Translational Modifications (PTMs) of proteins which influence their overall spatial 3D conformation, thereby affecting their stability, activity, and/or cellular localization (1). Reversible phosphorylation of serine, threonine or tyrosine residue has been the most extensively studied PTM (2). However, very often, phosphorylation of a protein is not solitary responsible to modulate its function. Rather, protein phosphorylation ensures interaction with its downstream protein-interacting partners which ultimately regulates its function. 14-3-3 proteins serve as prototype for such novel class of scaffold modules that recognize phosphor-serine/threonine (pS/pT) containing conserved binding motifs in a variety of signaling proteins. 14-3-3s are highly evolutionarily conserved dimeric (homo- and heterodimers), acidic proteins, widespread in almost all eukaryotic organisms (3–5). Although high degree of sequence conservation among 14-3-3 isoforms suggests a functional redundancy, the presence of phenotypes for single and multiple 14-3-3 knock-out mutants, and differential subcellular localization of 14-3-3s within a cell suggest that 14-3-3 isoforms *selectively* bind to their *individual* protein ligands with *different* affinities owing to the spatiotemporal regulation of the expression of different isoforms (6–10). Moreover, the large number of 14-3-3 isoforms expressed in an organism suggests high combinatorial complexity in dimer re-arrangement, which in turn fine-tunes their cellular functions.

14-3-3 proteins often interact with their cognate protein partners through canonical phosphorylated motifs, categorized as: Mode I (RXXpS/pT), Mode II (RXXXpS/pT) and Mode III (RXXpS/pTX_1-2_C’), where X is any amino acid and pS/pT represents phosphoserine or phosphothreonine (11–16). Mechanistically, 14-3-3 dimer interacts with its target protein(s) via two amphipathic grooves harbored by each monomer, and confers a conformational change that results in modulating functional parameters of its target protein(s) (17, 18). Depending on the biochemical nature of their phosphorylated protein targets, physical association with 14-3-3 proteins can have different functional consequences, resulting in modulation of its enzymatic activity, subcellular sequestration, protein stability and/or alteration of protein-protein interactions (9, 19). Earlier reports also suggest that, 14-3-3 possess chaperone-like activity akin to that of sHsps (small Heat shock proteins) and plays a critical role in formation of 14-3-3 mediated aggresomal targeting complex in response to accumulation of various misfolded proteins under conditions of cellular stress (20). Modulation in functional activity of kinases has also been attributed to 14-3-3 proteins, which have been described as inhibitors or activators of calcium and phospholipid dependent Protein Kinase C (PKC), and an activator of Raf-1 (21–24).

14-3-3 proteins, thus, operate as multipurpose *conformation (allosteric) clamps* that are recruited to hold its phosphorylated cognate protein(s) in place in response to cellular signaling pathways, culminating in regulation of apoptosis, adhesion-dependent integrin signaling, cell cycle control in response to genotoxic stress, ion-channels functioning, etc., thus governing diverse physiological processes and cellular status (4, 25– 28). In this regard, stability of 14-3-3 has been put forward as a basis for *“molecular anvil hypothesis”* according to which the rigid 14-3-3 dimer can induce structural rearrangements in its partner protein molecule(s), thereby regulating its functional properties while itself undergoing only minimal structural alterations (29). Studies on 14-3-3/client-protein interactions, by utilizing various biochemical and biophysical tools, may therefore provide tremendous opportunities for therapeutic interventions under various pathological conditions.

In the malaria parasite *P. falciparum*, two 14-3-3 isoforms have been annotated by database curators based on sequence similarity with experimentally annotated orthologs: *Pf*14-3-3I and *Pf*14-3-3II (accession numbers: PF3D7_0818200 and PF3D7_1362100, respectively), as documented in PlasmoDB Plasmodium Genomic Resource database (release 46; updated on 6th Nov., 2019). The findings of the present study confirm the presence of 14-3-3 protein in the malaria parasite. Further, we report that 14-3-3I interacts with a highly expressed protein in the parasite, Calcium-Dependent Protein Kinase 1 (CDPK1) that plays key role in multitude of essential cellular processes, including parasite invasion and egress during intra-erythrocytic proliferative stages of the parasite. Antagonizing the 14-3-3I/CDPK1 protein-protein interaction (PPI) by utilizing well characterized 14-3-3 recognition motifs as CDPK1 mimetic inhibits parasite growth *in vitro*, which indicates crucial physiological role of this PPI in the parasite. Our work sets a precedent for the rational design of 14-3-3 based PPI inhibitors by utilizing 14-3-3 recognition motif peptides, as a potential antimalarial strategy.

## Results

### Sequence analysis and identification of P. falciparum 14-3-3I

MultAlin-based sequence alignment of 14-3-3 isoforms from *Homo sapiens* and *P. falciparum* 3D7 demonstrated patterns of conservation and correlation in *Pf*14-3-3I protein sequence in light of the well-studied orthologs in humans (Figure 1A). Residues with high consensus value (>90%) are shaded in red and residues with low consensus value (>50%, <90%) are shaded in blue. α-helices and Nuclear Export Signal (NES) are also indicated. Five highly conserved sequence blocks, as identified by Wang W. and Shakes DC. (1996) were observed, as shown boxed and shaded red (30). Residues at the dimerization interface (solid circles) and residues involved in phosphopeptide target binding (solid squares) were found to be conserved in all 14-3-3 isotypes, except *Pf*14-3-3II which seems to be the most divergent form of 14-3-3 proteins. Based on literature review and comparative sequence analysis with the *Hs*14-3-3 isoforms, probable amino acid residues of *Pf*14-3-3I involved in dimerization and phosphopeptide (target) binding were identified and highlighted in the overall *Pf*14-3-3I architecture (Figure 1B).

**Figure 1:**
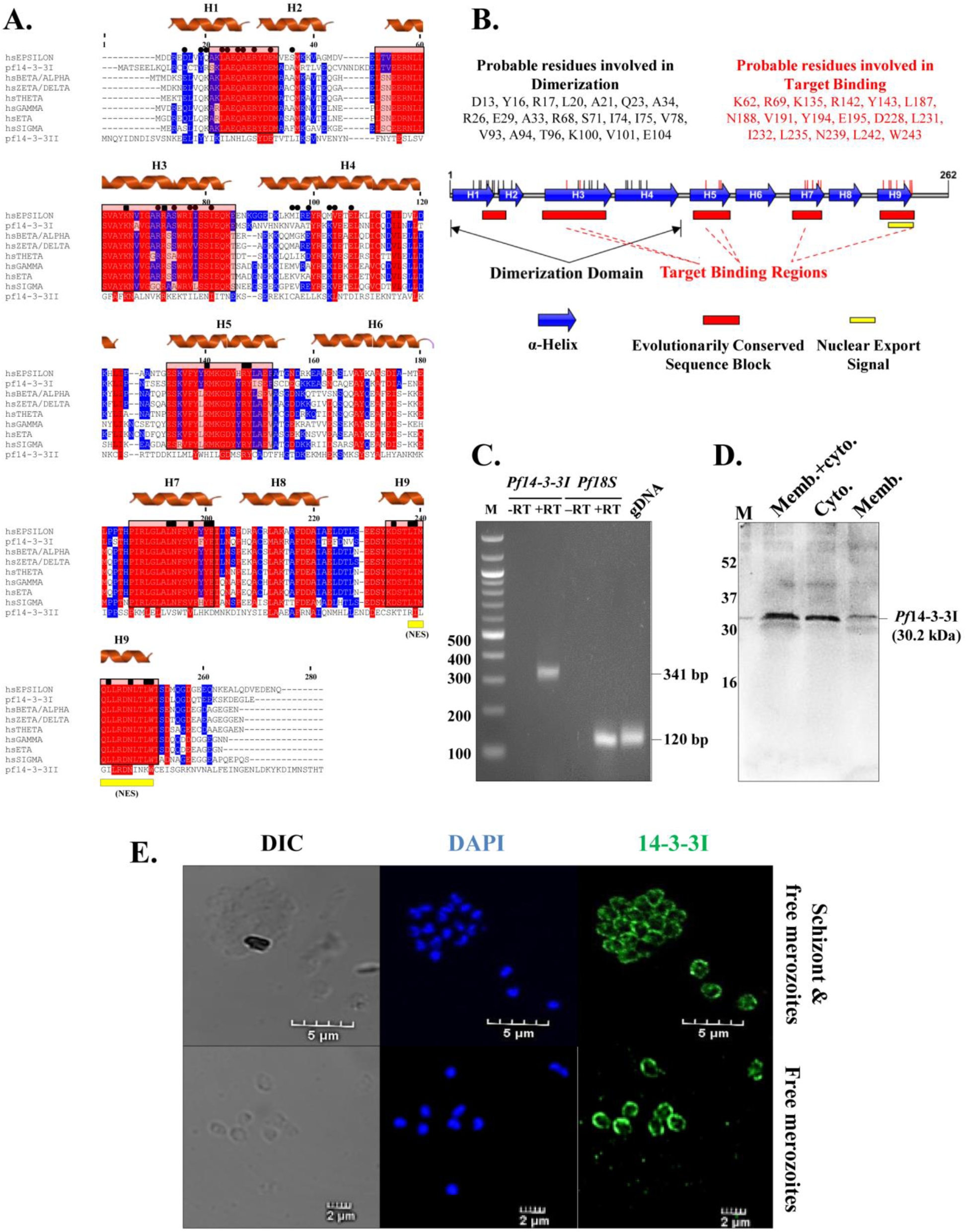
Comparative sequence analysis, protein architecture and identification of *P. falciparum* 14-3-3I. **A)** *MultAlin-based sequence alignment*. Multiple sequence alignment was performed to contextualize patterns of conservation and correlation in *Pf*14-3-3 protein sequences in light of well-studied orthologs in humans. Residues with high (>90%) and low (>50%, <90%) consensus values are shaded in red and blue, respectively. α-helices and NES are indicated. Five highly conserved sequence blocks are shown boxed and shaded red. Residues involved in dimerization (solid circles) and phosphopeptide binding (solid squares) are conserved in all 14-3-3 Isotypes, except *Pf*14-3-3II which appears to be the most divergent form. **B)** *Overall Pf14-3-3I architecture*. Based on multiple sequence alignment, probable amino acid residues of *Pf*14-3-3I involved in dimerization and target binding are shown. *Pf*14-3-3I architecture was constructed by using Illustrator for Biological Sequences (IBS 1.0.3). **C)** *Detection of 14-3-3 encoding cDNA. Pf*14-3-3I transcript was amplified from cDNA prepared from schizonts, by using *pf14-3-3I* specific primer sets. Desired band size of 341 bp was seen. Amplification of *pf18S* was taken as positive control. The experiment was done twice. **D)** *Western blot analysis of 14-3-3I*. In-house generated polyclonal sera raised in Balb/c mice against r14-3-3I was used to confirm the existence of native 14-3-3I protein in parasite (schizonts) lysate. Desired protein band of 30.2 kDa was observed in both cytosolic and membrane fractions of the lysate. The experiment was done twice. **E)** *Localization of 14-3-3*. Anti-r14-3-3I mice serum was used to probe localization of the protein in mature schizonts and free merozoites by using confocal microscopy. 14-3-3 protein was found to be localized towards cell periphery. The experiment was done thrice. H1-H9: α-helices; M: DNA or protein marker; RT: Reverse Transcriptase; DIC: Differential Interference Contrast image.

X-Ray diffraction based structural model of *Hs*14-3-3 epsilon_dimer_ was used as a template to generate three-dimensional coordinates of *Pf*14-3-3I_dimer_. After optimal rigid-body superimposition of *Hs*14-3-3 epsilon_dimer_ with the generated structural model of *Pf*14-3-3I_dimer_, overall Root-Mean-Square Deviation (RMSD) value of the C-alpha atomic coordinates was found to be 0.63 Å, suggesting a reliable 3D structure of *Pf*14-3-3I_dimer_. Structural model of *Pf*14-3-3I_dimer_ revealed strong resemblance with it’s counterparts in other living organisms, with overall folds forming a clamp like structure where each monomer is capable of forming a functional amphipathic groove for binding to phosphorylated residues on target proteins. Additionally, both monomers of *Pf*14-3-3I_dimer_ were found to be oriented in opposite direction with respect to each other. Helical regions in one of the monomers are marked from H1 to H9 (Supporting figure S1A). Assessment of stereochemical quality and accuracy of the generated homology model displayed 88.7% of amino acid residues lying in the most favored (“core”) regions, with 8.8%, 1.5%, and 1.1% residues in “additional allowed”, “generously allowed” and “disallowed regions” of Ramachandran plot, respectively (Supporting figure S1B). Since, protein structure with ≥90% of its amino acid residues lying in the most favoured regions of Ramachandran plot is considered to be as accurate as a crystal structure at 2Å-resolution, this indicated that the backbone dihedral angles: phi and psi of the generated *Pf*14-3-3I_dimer_ model were reasonably accurate. The comparable Ramachandran plot characteristics and RMSD value confirmed the reliability of the 3D-structural model of *Pf*14-3-3I_dimer_ to be taken further for docking and simulation studies.

Unrooted phylogenetic relationship of *Pf*14-3-3 isoforms with their orthologs present across three major kingdom of life: plantae, animalia and fungi showed that *Pf*14-3-3 isoforms have followed convergent evolutionary pathway with 14-3-3 proteins from plant *non-epsilon* group (Supporting figure S2A). Branches with green, red and blue squares belong to kingdom plantae, animalia and fungi, respectively. Detailed evolutionary relationship of *Pf*14-3-3 isoforms with their orthologs from plant *non-epsilon* group is shown (Supporting figure S2B).

Transcripts encoding for 14-3-3I protein was amplified from cDNA prepared from merozoites, by using *pf14-3-3I* specific primers, as mentioned in experimental procedures section. Detection of desired DNA fragment of 341 bp confirmed the existence of *14-3-3I* encoding cDNA in the parasite (Figure 1C). Amplification of transcripts encoding for 18S rRNA (120 bp) by using *pf18s* specific primers was taken as positive control. Further, western blot analysis of native 14-3-3I protein in the parasite (schizonts) lysate, by using in-house generated polyclonal mice sera raised against r14-3-3I, confirmed the existence of 14-3-3I protein in the parasite (Figure 1D). Desired protein band of around 30.2 kDa was observed in both Cytosolic (C) and Membrane (M) fractions of the parasite lysate. IFA in mature schizonts and free merozoites with the anti-r14-3-3I mice serum indicated localization of the protein towards cell periphery.

To update optimal 14-3-3 binding consensus motifs, literature search followed by mining of publically available databases was done to identify all experimentally validated 14-3-3 interacting partners present in prokaryotes & eukaryotes, and collate details of phosphoSer/Thr target sites on the target proteins. In total, 323 mode I sites from 243 target proteins, 81 mode II sites from 77 target proteins and 9 mode III sites from 9 target proteins were identified as gold-standard 14-3-3 binding phosphoSer/Thr sites (Supporting table S1). *Updated* optimal consensus 14-3-3 binding motifs constructed from amino acid sequences of these 14-3-3 binding phosphopeptides are shown (Supporting Figure S3A). Further, protein kinases were filtered out as putative binding partners of *Pf*14-3-3I by combining publicly available global phospho-proteomic datasets of peptides enriched from schizont stage of *P. falciparum*, with 14-3-3 binding sequence motifs search (Supporting Figure S3B). Our results speculate that *Pf*14-3-3 binds to and regulates the physiological activity of master kinases of the parasite: CDPK1, Protein Kinase G (PKG), Protein Kinase A regulatory subunit (PKA_R_) and Protein Kinase A catalytic subunit (PKA_C_). Amino acid sequences of probable 14-3-3 binding phosphopeptides of CDPK1 are also shown (Supporting Figure S3A).

### Molecular Dynamics (MD) Simulation reveals stable complex formation of 14-3-3I_dimer_ with CDPK1

To probe the associated molecular interactions that regulate the binding affinity of 14-3-3I_dimer_ with *p*CDPK1, we performed MD simulations. A schematic representation of the interactions between 14-3-3I_dimer_ and *p*CDPK1 is shown in Figure 2A. The phosphorylated serine, *p*S64 of pCDPK1 was found to play a key role in mediating its interaction with 14-3-3I_dimer,_ where a Hydrogen-bond was formed between *p*S64 of pCDPK1 and K227 of 14-3-3I_dimer_ with a bond length of 2.92Å. In addition to Hydrogen-bonding, the 14-3-3I_dimer_-*p*CDPK1 binding was further stabilized by hydrophobic interactions between *p*S64 of *p*CDPK1 and Y226, Y555, E557, I597 of 14-3-3I_dimer_ located within the amphipathic binding pocket of the protein. Further comprehensive analysis of the MD trajectories, *i*.*e*., Root Mean Square Deviation (RMSD), Radius of Gyration and variation in the Hydrogen-bond formation between 14-3-3I_dimer_ and *p*CDPK1 were also reported as a function of simulation time (Figure 2A). The variation of RMSD and Radius of Gyration in 14-3-3I_dimer_ and *p*CDPK1 clearly indicated a lesser stability and compactness in the former compared to *p*CDPK1. The higher conformational fluctuations in 14-3-3I_dimer_ may assist in forming a stable complex with *p*CDPK1 via induced-fit mechanism (31, 32). The higher stability of the protein-protein complex was also confirmed by the extensive Hydrogen-bond formation seen between *p*CDPK1 and 14-3-3I_dimer_ at different time intervals throughout the simulation and a negative binding energy (−36.8 ± 1.0 kcal/mol) as obtained from MM/GBSA method. Movie S1 shows the interactions between 14-3-3_dimer_ (blue) and *p*CDPK1 (grey).

**Figure 2:**
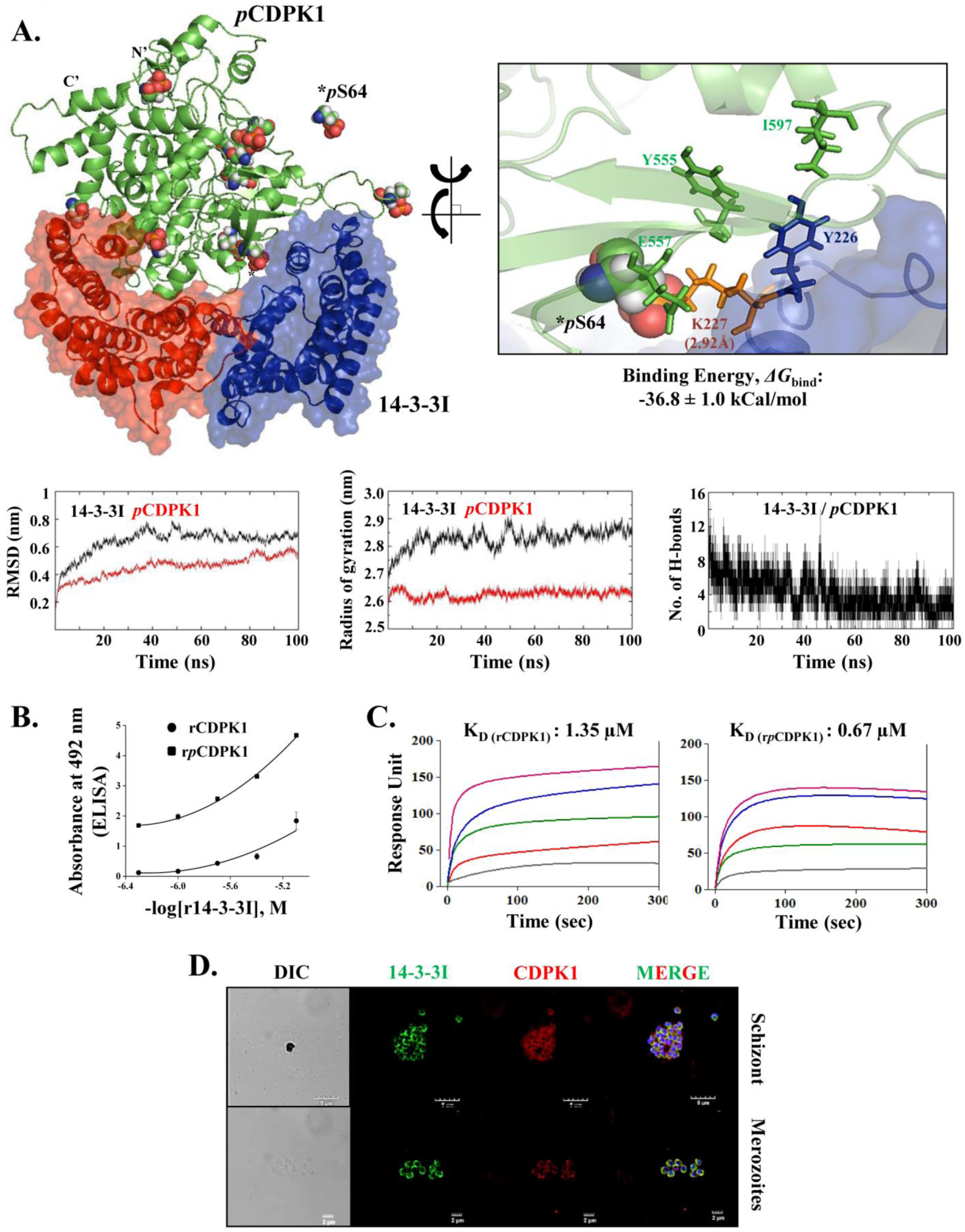
*Pf*14-3-3I interacts with *Pf*CDPK1. **A)** *Schematic representation of the interactions between pCDPK1 and 14-3-3I*_*dimer*_. A Hydrogen-bond formation is shown between *p*S64 of *p*CDPK1 and K227 of 14-3-3I_dimer_ which has a bond length of 2.92Å. The variation in RMSD and Radius of Gyration of 14-3-3I_dimer_ & CDPK1 is shown as a function of simulation time. The variation in number of intermolecular Hydrogen-bond formation between 14-3-3I_dimer_ and CDPK1 is shown as a function of simulation time. **B)** *ELISA*. For 14-3-3I/CDPK1 interaction analysis, a kinase reaction was set up with rCDPK1 (100 ng/µl), in the absence & presence of Ca^2+^-ions for conditions requiring rCDPK1 and r*p*CDPK1, respectively. rCDPK1 or r*p*CDPK1 (200 ng) was coated onto Poly-L-Lysine coated microtitre plates, followed by incubation with different concentrations of r14-3-3I (0.5 to 8 μM). Interaction analysis was done by using monoclonal antibody against GST protein. r14-3-3 was found to interact with rCDPK1 in a phosphorylation dependent manner. The experiment was done twice in triplicates. **(C)** *Concentration dependent real-time sensograms for SPR based biomolecular interaction analysis*. ELISA based 14-3-3I/CDPK1 interaction analysis was further confirmed with AutoLab Esprit SPR. rCDPK1 & r*p*CDPK1 showed differential binding affinities for r14-3-3I, with K_D_ values varying from 0.67 ± 0.0036 μM and 1.35 ± 0.0083 μM, respectively. The experiment was done twice. **(D)** *Co-localization of 14-3-3I and CDPK1*. 14-3-3I was found to co-localize very nicely with CDPK1 towards cell periphery in mature schizonts and free merozoites. Images were acquired by using Nikon A1-R confocal microscope. The experiment was done thrice. RMSD: Root Mean Square Deviation; pS: Phosphorylated serine; K_D_: Affinity constant; DIC: Differential Interference Contrast image.

### 14-3-3I exhibits divergent binding affinities for CDPK1 and pCDPK1 in vitro

For ELISA based 14-3-3/CDPK1 PPI analysis, Poly-L-Lysine coated 96-welled microtitre plate was coated with rCDPK1 (or r*p*CDPK1), followed by addition of increasing concentrations of the prey protein, r14-3-3I. The interaction analysis was done by using monoclonal antibody against GST protein. A concentration dependent binding between rCDPK1 (or r*p*CDPK1) and the prey protein r14-3-3I was observed (Figure 2B). PPI analysis by ELISA suggested that phosphorylation status of CDPK1 dictates its interaction with 14-3-3I. To quantitate the interaction between rCDPK1 (or r*p*CDPK1) and r14-3-3I, SPR analysis was performed by utilizing AutoLab Esprit SPR. r14-3-3I was immobilized at an average density of 4.3 ng per 1 mm^2^ of the sensor chip surface. Once immobilized, r14-3-3I demonstrated good stability throughout the experiment. Interaction analysis was done by injecting serial dilutions of either rCDPK1 or r*p*CDPK1 ranging from 100 nM to 1 μM over the r14-3-3I-immobilized sensor chip surface, followed by comparing their respective kinetics & binding affinities at RT. With increase in mass concentration of rCDPK1 and r*p*CDPK1, gradual increase in sensor signal was observed which linearly correlated with corresponding change in refractive index of the medium immediately adjacent to the SPR sensing surface. The concentration dependent real-time sensorgrams along with K_D_ values of interactions are shown in figure 2B. rCDPK1 and r*p*CDPK1 showed differential binding affinities r14-3-3I, with K_D_ values varying from 0.67 ± 0.0036 μM (14-3-3I/*p*CDPK1) and 1.35 ± 0.0083 μM (14-3-3I/CDPK1). Also, SPR sensograms suggested specificity of CDPK1 towards 14-3-3I cavity(ies) in terms of shape complementarity and chemical functionality.

IFA was performed on synchronized *P. falciparum* 3D7 culture to check for co-localization of 14-3-3I and CDPK1 by probing mature stages of the parasite with anti-*Pf*14-3-3I mouse-serum and anti-*Pf*CDPK1 rabbit-serum, and images were acquired using Nikon A1-R confocal microscope using the NIS Elements software. 14-3-3I protein was found to co-localize very nicely with CDPK1 protein towards cell periphery in mature schizonts and free merozoites.

### MD Simulation reveals stable complex formation of 14-3-3I_dimer_ with Phosphopeptides 1, 2

In Figure 3A, the interactions of 14-3-3I_dimer_ with phosphopeptides 1 and 2 were studied by performing 20 ns MD simulations. The peptide 1 was found to bind in the amphipathic groove of the receptor protein, 14-3-3I_dimer_. Here, three hydrogen bonds were formed between the phosphorylated Serine in peptide 1 and K306, K379, S302 of the receptor protein, in addition to hydrophobic interactions that further stabilized the 14-3-3I_dimer_-peptide complex. Similarly, the phosphorylated Serine of peptide 2 formed a Hydrogen-bond with N188 of 14-3-3I_dimer_, along with stabilizing hydrophobic interactions. A comparative study using time-dependent variations in RMSD and Radius of Gyration, however, revealed that 14-3-3I_dimer_-peptide 2 complex was more stable and slightly more compact compared to when 14-3-3I_dimer_ interacts with the peptide 1. The observation was supported by the higher number of intermolecular Hydrogen-bonds that are formed between 14-3-3I_dimer_ and peptide 2 as compared to peptide 1. This also accounts for the higher negative binding energy observed for 14-3-3I_dimer_-peptide 2 complex (−41.8 ± 1.0 kcal/mol) relative to 14-3-3I_dimer_-peptide 1 complex (−67.2 ± 0.1 kcal/mol). Movies S2 and S3 show the interactions of 14-3-3_dimer_ (blue) with peptides 1 & 2 (yellow), respectively.

**Figure 3:**
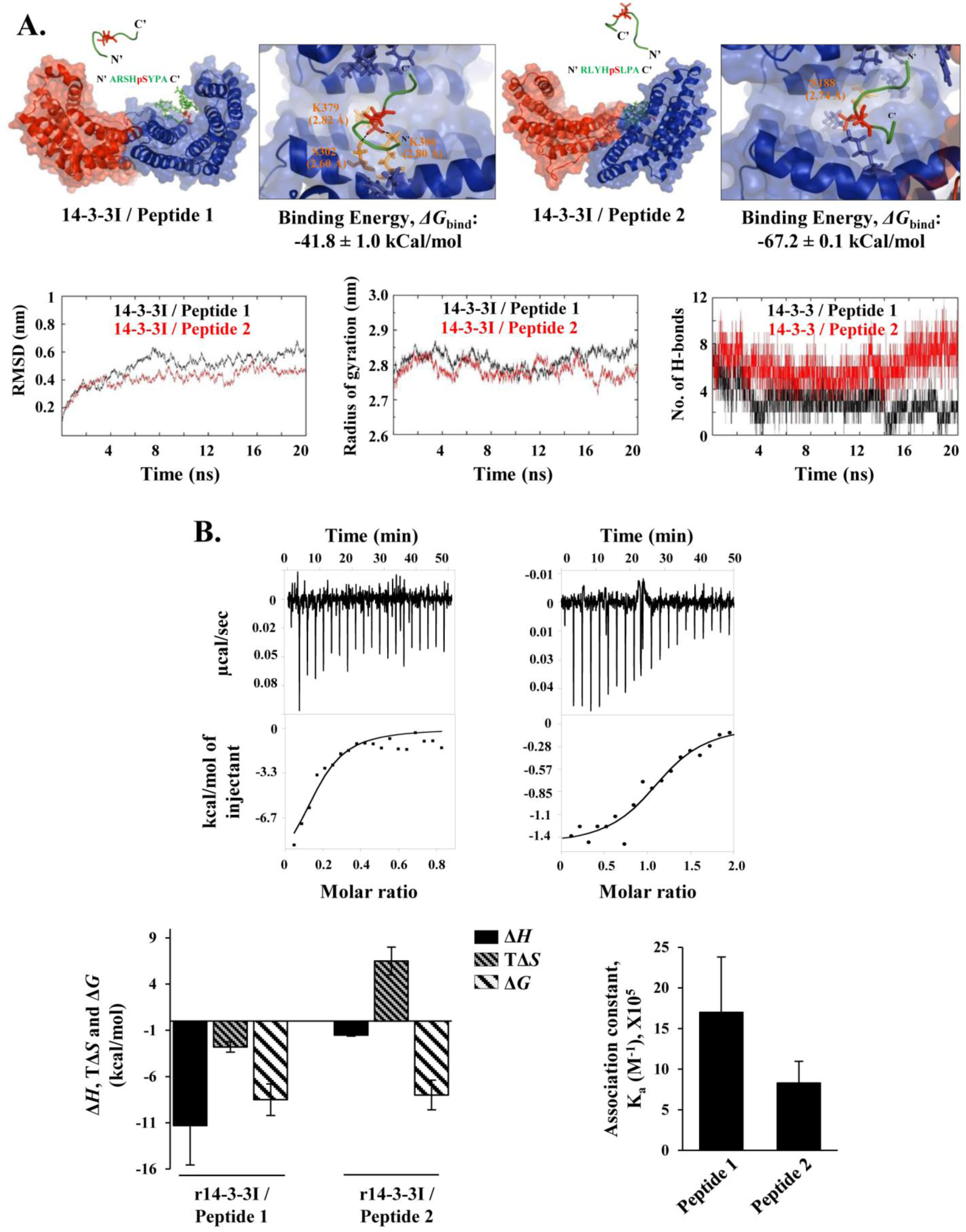
Phosphopeptides 1 & 2 interact with *Pf*14-3-3I. **A)** *Schematic representation of the interactions between 14-3-3I*_*dimer*_ *and phosphopeptides 1 (left) & 2 (right)*. The Hydrogen-bonds formed between phosphorylated Serine of peptide 1 and K306, K379, S302 of 14-3-3I_dimer_ (left); and, phosphorylated Serine of peptide 2 with N188 of 14-3-3I_dimer_ (right). The variation in RMSD and Radius of Gyration of 14-3-3I_dimer_ when it interacts with peptides 1 & 2 as a function of simulation time. The variation in number of intermolecular Hydrogen-bond formation between 14-3-3I_dimer_ and peptides 1 & 2 as a function of simulation time. **B)** *Representative binding isotherms resulting from titrations of r14-3-3I with phosphopeptides 1 & 2*. ITC was employed to determine kinetic parameters of complexation between r14-3-3I (10 μM) and peptide 1 (40 μM) or 2 (100 μM), by using MicroCal iTC200. On the basis of K_a_ values, binding strength was found to be higher in case of 14-3-3I/peptide 1 (K_a_: 1.7 × 10^6^ ± 6.8 × 10^5^ M^−1^) than in case of 14-3-3I/peptide 2 (K_a_: 8.3 × 10^5^ ± 2.67 × 10^5^ M^−1^) interaction. The binding of peptide 1 with r14-3-3I was enthalpically favorable whereas entropically unfavourable, resulting in strong binding free energy. For 14-3-3I/peptide 2 complex, the interaction was enthalpically as well as entropically driven. Experimental data is represented as the amount of heat released per second (μcal/sec; corrected for heat of dilution of the ligand) following each injection of the ligand (peptide) into the receptor protein (r14-3-3I), as a function of time (min.). The data were fitted by using single-site binding model. Solid line represents the best fit of the non-linear experimental data. The experiment was done twice. RMSD: Root Mean Square Deviation; Δ*H*: change in Enthalpy; Δ*S*: change in Entropy; Δ*G*: change in Gibbs free energy; T: Temperature.

### Phosphopeptides 1 & 2 interact with 14-3-3I in vitro

After affirming 14-3-3I_dimer_ interaction with phosphopeptides 1 & 2 through molecular dynamics simulation studies, we further sought to establish the knowledge of binding modes of complexation between r14-3-3I and the peptides, by employing ITC. This was achieved by determining kinetic parameters of the interaction, like binding affinity constant (K_a_) and change in enthalpy (Δ*H*), under sustained salt (150 mM NaCl) and pH (7.4) conditions, at RT. Δ*H* along with K_a_ values were then utilized for calculation of additional parameters, like binding free energy (Δ*G*, equivalent to −RT.lnK_a_) and entropy [Δ*S*, equivalent to (Δ*H* - Δ*G*)/T]. The representative binding isotherms resulting from titration of peptide 1 or 2 with r14-3-3I are represented in figure 3B. Binding isotherm with peptide 1 was monophasic in nature, reaching a plateau phase indicating saturation of r14-3-3I binding sites. In case of peptide 2, as further injections continued, decline in exothermic heat resulted in sigmoidal curve ending near zero baseline. On the basis of the nature of the curves, the data were fitted by using single-site binding model. In Figure 3B, solid line shows the best fit of non-linear experimental data, and the model reproduces experimental data fairly well. On the basis of K_a_ values, binding strength was found to be higher in case of 14-3-3I/peptide 1 (K_a_: 1.7 × 10^6^ ± 6.8 × 10^5^ M^−1^) than in case of 14-3-3I/peptide 2 (K_a_: 8.3 × 10^5^ ± 2.67 × 10^5^ M^−1^). Moreover, it was observed that the binding of peptide 1 with r14-3-3I protein was enthalpically favorable (Δ*H* = −11.34 ± 4.27 kcal/mol), whereas entropically unfavourable (TΔ*S* = −2.8 kcal/mol (± 15-20%), resulting in strong binding free energy (Δ*G* = −8.5 kcal/mol (± 15-20%). For 14-3-3I/peptide 2 complex, the interaction was enthalpically as well as entropically driven (Δ*H* = −1.52 ± 0.096 kcal/mol, TΔ*S* = 6.5 kcal/mol (± 15-20%) and Δ*G* = −8.0 kcal/mol (± 15-20%).

### Phosphopeptides 1 & 2 as CDPK1 mimetic to antagonize 14-3-3I/CDPK1 interaction

ELISA based 14-3-3/CDPK1 PPI analysis was performed in the presence of varying concentrations of phosphopeptides 1 & 2. The interaction analysis was done by using monoclonal antibody against GST protein. Concentration dependent inhibition of binding between r*p*CDPK1 and r14-3-3I was observed (Figure 4A). To further confirm 14-3-3I/CDPK1 interaction inhibition by the peptides, western blot based GST pull-down assay was performed in which r14-3-3I (or GST, as negative control) was coupled with Glutathione Sepharose® 4B beads, followed by binding with rCDPK1 or r*p*CDPK1 to form bead-bound protein complexes, in the absence and presence of 10 μM concentration of the peptides. Immunoblotting with HRP-conjugated anti-His antibody indicated that phosphorylation status of CDPK1 dictates its interaction with 14-3-3I, as confirmed by ELISA (Figure 4B). Moreover, inhibition of binding between rCDPK1 (or r*p*CDPK1) and r14-3-3I was readily observed in the presence of peptides 1 & 2. The same blot stripped and re-probed with HRP-conjugated anti-GST antibody served as experimental control to check equal coupling of r14-3-3I (or GST) with Glutathione Sepharose beads in all binding reactions.

**Figure 4:**
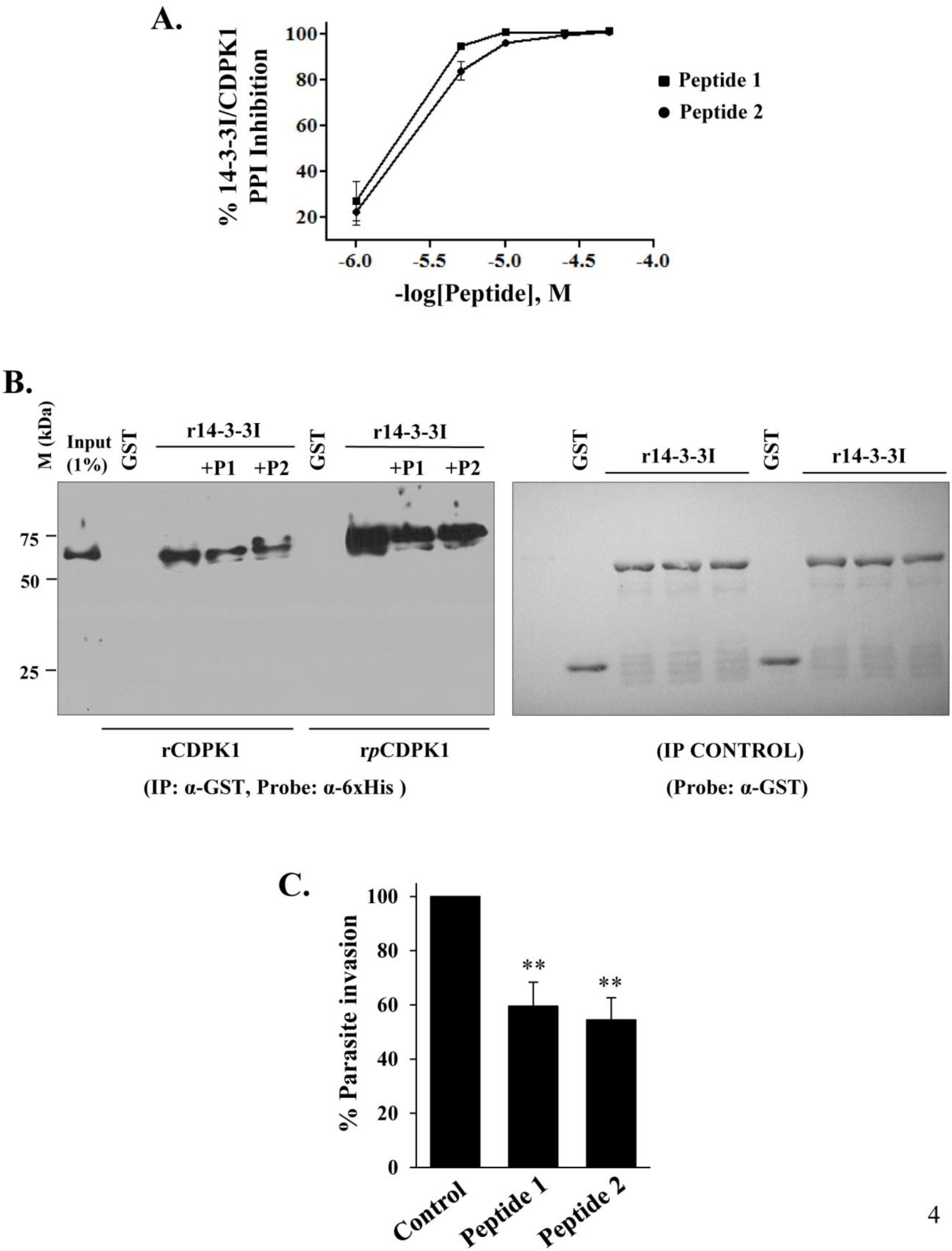
Phosphopeptides 1 & 2 as CDPK1 mimetic to antagonize *Pf*14-3-3I/*Pf*CDPK1 interaction. **A)** *ELISA based interaction inhibition*. Concentration dependent inhibition in binding of r*p*CDPK1 with r14-3-3I was observed in the presence of phosphopeptides 1 & 2 (1 μM to 50 μM). Interaction analysis was done by using monoclonal antibody against GST protein. The experiment was done twice in triplicates. **B)** *GST-based pull-down assay*. To further confirm 14-3-3I/CDPK1 interaction inhibition, western blot based GST pull-down assay was performed in which Glutathione Sepharose® 4B beads was coupled with r14-3-3I (10 µg), followed by incubation with rCDPK1 or r*p*CDPK1 (10 µg), in the absence and presence of peptides 1 & 2 (10 μM). Immunoblotting with HRP-conjugated anti-His antibody indicated that phosphorylation status of CDPK1 dictates its interaction with 14-3-3I. Moreover, interaction inhibition was readily observed in the presence of both peptides. Same blot stripped and re-probed with HRP-conjugated anti-GST antibody confirmed equal coupling of r14-3-3I (or GST) with Glutathione Sepharose beads in all binding reactions. The blots shown are representative of two independent experiments. **C)** *Parasite invasion assay*. Mature schizonts were allowed to invade into erythrocytes in the presence of peptides 1 or 2 (12.5 μM). % parasite growth inhibition was calculated by FACS using BD LSRFortessa™ cell analyzer. Untreated parasites served as control. Both peptides significantly inhibited progression of schizonts to ring stage. Control parasites were healthy, producing rings at 24 hrs post-incubation. The experiment was done twice in triplicates. M: Protein marker; P1: Phosphopeptide 1; P2: Phosphopeptide 2.

Phosphopeptides 1 & 2 with propensity to bind 14-3-3I protein and inhibit 14-3-3I/CDPK1 interaction, were subjected to *P. falciparum* growth inhibition *in vitro*. Towards this, mature (punctated) schizonts were allowed to invade into erythrocytes in the presence of 12.5 μM concentration of the peptides. Untreated parasites served as control and percentage parasite growth inhibition was measured by flow cytometry, as described in experimental procedures section. Peptides 1 & 2 showed significant inhibition in progression of schizonts to ring stage of the parasite (Figure 4C). Control parasites were healthy, producing rings at 24 hrs post-incubation.

## Discussion

14-3-3 is a novel class of dimeric, conserved scaffold proteins that recognize phosphor-serine/threonine (pS/pT) containing conserved binding motifs in a variety of signaling proteins, thus regulating their physiological functions. Whereas yeast has two genetically distinct but structurally homologous isoforms, and mammals seven, most plant genomes contain a dozen 14-3-3 genes which can be divided into two distinct groups: epsilon and non-epsilon, based on their sequence homology and exon/intron structure (3, 4, 33, 34). The epsilon group of 14-3-3 proteins is considered as *ancestral* and destined to fulfill fundamental cellular functions. Contrastingly, the non-epsilon group evolved later and members of this group might be related to organism *specific* functions (35). The large number of 14-3-3 isoforms expressed in an organism suggests high combinatorial complexity in dimer re-arrangement, thus fine-tuning their cellular functions.

There are only a few evidences in literature which supports the existence of 14-3-3 protein in Plasmodium species. Identification of gene encoding 14-3-3 protein in *P. falciparum* and *P. knowlesi* was first reported by Basima Al-Khedery *et al* (1999), wherein authors reported that *Pk*14-3-3 transcript begins to be expressed in the ring-stage, predominates in young trophozoites, and thereafter declines during asexual intra-erythrocytic proliferative stages of the parasite. Also, antiserum produced against recombinant *Pk*14-3-3 was able to cross-react with *Pf*14-3-3 (36). Later, Marco Lalle *et al* (2011) showed that in *P. berghei* parasitized erythrocytes, the host cytoskeletal protein demantin is repositioned to the parasite where it interacts with *Pb*14-3-3 in a phosphorylation dependent manner, thus influencing the remodeling of the erythrocytic cytoskeleton and modulating the host erythrocyte invasion (37). Further, Eeshita G. Dastidar *et al* (2013) identified *Pf*14-3-3 as a novel member of *P. falciparum* histone phosphosite binding protein repertoire, isolated from asexual blood stages of the parasite (38).

14-3-3 isoforms are also annotated in *P. falciparum* by database curators: *Pf*14-3-3I and *Pf*14-3-3II (accession numbers: PF3D7_0818200 and PF3D7_1362100, respectively), as documented in PlasmoDB Plasmodium Genomic Resource database (release 46; updated on 6th Nov., 2019). Upon comparative sequence analysis with structurally and functionally characterized orthologs from *Homo sapiens*, we report patterns of sequence and structural conservation in *Pf*14-3-3I protein sequence particularly encompassing residues at the dimerization interface and those involved in phosphopeptide binding (Figure 1A, 1B, S1). Concomitantly, our phylogenetic analysis shows that despite having evolved separately since the early eukaryotes, *Pf*14-3-3I protein has followed convergent evolution with plant *non-epsilon* group, as depicted in the phylogenetic model (Figure S2). We further confirmed the presence of 14-3-3 protein in the malaria parasite by utilizing various molecular biology techniques. Real-time PCR indicated presence of transcript encoding for *Pf*14-3-3I protein (Figure 1C). Presence of full length protein was confirmed by western blot analysis in the parasite lysate, by using in-house generated polyclonal sera raised in mice against recombinant *Pf*14-3-3I protein (Figure 1D). Confocal microscopic imaging in mature schizonts and free merozoites indicated that *Pf*14-3-3I protein localizes at the cell periphery (Figure 1E).

14-3-3 dimer acts as multipurpose ‘allosteric clamp’ which holds and confers conformational changes in its cognate phosphorylated protein(s) (39). In this mode of interaction, docking site for 14-3-3 is created by a combination of inherent shape and phosphate added in the target protein in response to a particular signaling cascade. Association with 14-3-3 renders modulation in enzymatic activity, subcellular localization and/or stability of the target protein(s) (4, 9, 19, 25–28, 40). Kinases have also been reported to be functionally modulated as a consequence of their interaction with 14-3-3 proteins. In this regard, first report came from Toker A. *et al* (1990) who identified 14-3-3 as one of the potent inhibitors of calcium dependent Protein Kinase C (PKC) in sheep brain, where it could inhibit phosphorylation activity of the kinase over a wide range of calcium-ion concentrations (21). In contradiction to this study, Tanji M. *et al* (1994) identified 14-3-3 isoforms as activators of PKC in bovine forebrain (23). Unlike previously reported, the activation of PKC by 14-3-3 was found to be independent of phosphatidylserine and calcium-ions and as such, provided an alternative mechanism for the activation of PKC that obviates its translocation to membranes. In the concurrent year, Wendy J. Fanti *et al* (1994) reported that 14-3-3 isoforms bind and enhance serine/threonine kinase activity of Raf-1, a key mediator of mitogenesis and differentiation, thus promoting Raf-1 dependent oocyte maturation (24). Later, Acs P. *et al* (1995) reported differential activation of various PKC isoforms, especially PKC epsilon, by 14-3-3 zeta protein (22).

In the current study, we report that 14-3-3I interacts with Calcium-Dependent Protein Kinase 1 (CDPK1), a highly expressed protein in the malaria parasite, *P. falciparum* that plays key role in multitude of essential cellular processes, including parasite invasion and egress during intra-erythrocytic proliferative stages of the parasite. Preliminary structural modeling and molecular dynamics simulation of 14-3-3I_dimer_/*p*CDPK1 complex indicated that CDPK1 binds favorably to amphipathic groove of the receptor protein. The interaction is driven by charged residues present at the interface of the two proteins, and hydrophobic interactions stabilize the association. The Gibb’s free energy for 14-3-3I_dimer_/*p*CDPK1 interaction, as calculated using Molecular Mechanics Generalized Born and Surface Area continuum solvation (MM/GBSA) method was found to be, Δ*G*_bind_: −36.78 ± 0.96 kcal/mol (Figure 2A) (41). Moreover, the structural changes conferred in 14-3-3I_dimer_ upon the binding of CDPK1 might assist a stable complex formation between them. The simulation results for 14-3-3I/CDPK1 interaction were confirmed experimentally by utilizing various biochemical and biophysical tools including ELISA, Surface Plasmon Resonance (SPR) and pull-down experiments with recombinant purified proteins, which indicated that the interaction is dependent on phosphorylation status of CDPK1 (Figure 2B, 2C, 4C). SPR analysis indicated that 14-3-3I has approximately two fold higher affinity towards phosphorylated CDPK1, with binding affinity constants (K_D_) of 1.35 µM & 0.67 µM for 14-3-3I/CDPK1 and 14-3-3I/*p*CDPK1 interaction, respectively (Figure 2C). In a related study by our group, the phosphorylation dependent interaction of 14-3-3I with CDPK1 was validated in merozoites, wherein, by pulling down 14-3-3I, CDPK1 was detected in a Ca^2+^ dependent manner (42). Our findings overlap with the conception that 14-3-3 proteins confer phosphorylation-dependent binding to their protein ligands. Further, confocal microscopic imaging in mature schizonts and free merozoites indicated that 14-3-3I co-localizes well with CDPK1 towards the cell periphery (Figure 2D).

Earlier reports suggest that synthetic peptides containing phosphorylated 14-3-3 binding motifs can efficiently inhibit the association of 14-3-3 proteins with their interacting partners, by typically binding to the conserved amphipathic groove of 14-3-3s. The first report utilizing peptides as 14-3-3 binding high-affinity antagonists came from Wang B. *et al* (1999), who identified unphosphorylated R18 peptide (PHCVPRDLSWLDLEANMCLP) by screening phage display libraries (43). The peptide exhibited high affinity for different isoforms of 14-3-3 by binding to the amphipathic groove of 14-3-3 via polar and hydrophobic interactions. R18 efficiently blocked the binding of 14-3-3 to Raf-1 kinase, a physiological ligand of 14-3-3, thereby effectively abolishing the protective role of 14-3-3 against phosphatase-induced inactivation of Raf-1, and virulence factor exoenzyme S (ExoS) of the pathogenic bacterium *Pseudomonas aeruginosa* (44). To further enhance R18 activity in cells, a dimeric R18 sequence, termed difopein (*di*meric *fo*urteen-three-three *pe*ptide *in*hibitor) was used to dissect the physiological role of 14-3-3/client protein interaction, and served as a basis for targeting pro-survival function of 14-3-3 as a potential anticancer strategy (45). Thereafter, other peptide-based inhibitors have been developed, such as the macrocyclic cross-linked peptide synthesized by Glas A. *et al* (2014) and Philipp MC. *et al* (2016) that inhibit the interaction between human 14-3-3 and ExoS (46, 47). The so-called ESp peptide (^420^QGLLDALDLAS^430^), a 14-3-3 binding motif of exoenzyme S, was used as starting point for the design of the macrocyclic PPI inhibitor with different cross-link architectures. Later, Milroy LG. *et al* (2015) synthesized modified Tau peptide ‘hybrids’ as inhibitor of 14-3-3/Tau interaction, a potential drug target for the treatment of Alzheimer’s disease (48). The peptides were designed with an extended hydrophobic area at the C-terminus that targeted the highly conserved pocket within the amphipathic groove of 14-3-3.

Based on the literature review and owing to the fact that well characterized 14-3-3 recognition motifs (Mode I, II and III) and other naturally occurring 14-3-3-binding peptides can be utilized as 14-3-3/client protein PPI antagonists, we utilized two different phosphopeptides: Mode I (ARSH*pS*YPA or *peptide 1*) and Mode II (RLYH*pS*LPA or *peptide 2*) (*pS*: phosphoserine) as CDPK1 mimetics to inhibit 14-3-3/CDPK1 PPI inhibition. These phosphopeptides were identified by Rittinger K. *et al* (1999) from oriented peptide library screening and are recognized by all mammalian and *S. cerevisiae* 14-3-3 isotypes (49). Preliminary structural modeling and molecular dynamics simulation of 14-3-3I_dimer_/peptide 1 or 2 complexes indicated that the peptides bind favorably to amphipathic groove of the receptor protein. The interaction is driven by Hydrogen-bonds formed between phosphorylated serine of peptide ligands and receptor protein, and hydrophobic interactions stabilize the association. The Gibb’s free energy for 14-3-3I_dimer_/peptide 1 and 14-3-3I_dimer_/peptide 2 interactions, as calculated using MM/GBSA method was found to be, Δ*G*_bind_: −41.84 ± 0.96 and −67.18 ± 0.06 kcal/mol, respectively (Figure 3A). The stable complex formation between 14-3-3I_dimer_ and phosphopeptides 1 & 2 confirms their effectiveness as potential inhibitors of 14-3-3I_dimer_ with its interacting partners. The simulation results for 14-3-3I/peptides interaction were confirmed experimentally by using Isothermal Titration Calorimetry (ITC) with recombinantly purified 14-3-3I protein and peptides synthesized *de novo*. Peptide 1 shows stronger binding affinity for r14-3-3I than peptide 2 (Figure 3B). Much stronger Δ*H* value for peptide 1 indicates its ability to form a strong network of Hydrogen-bonds and Van Der Waals interactions with 14-3-3I protein. The unfavorable entropy may be the result of loss of conformational freedom upon 14-3-3I/peptide 1 complex formation. In case of peptide 2, interactions are strongly dominated by favorable entropy, with a slight contribution from enthalpy. Positive entropy suggests important contribution from *solvation entropy*, which results from the loss of water molecules from the binding surfaces of interacting partners when they come together to form a complex (50). Small negative Δ*H* value indicates smaller number of H-bonding interactions contributing to the formation of 14-3-3I/peptide 2 complex. Further, both peptides 1 and 2 efficiently blocked the interaction of 14-3-3I with CDPK1 protein, as confirmed by ELISA and pull-down experiments with recombinantly purified proteins (Figure 4A and 4B). To dissect the physiological role of 14-3-3I/CDPK1 interaction, mature *P. falciparum* schizonts were allowed to rupture and invade into new erythrocytes, in the absence and presence of peptides 1 & 2. Both peptides significantly inhibited the progression of schizonts to ring stage (Figure 4C).

In conclusion, our findings confirm the existence of *Pf*14-3-3I protein in the malaria parasite *P. falciparum*, and present insight into its sequence and structural features which may prove to be an initial lead in understanding of its function in the parasite. Moreover, we have shown the interaction of *Pf*14-3-3I with CDPK1, an important kinase of the parasite involved in motility and apical organelle discharge critical for invasion process. This study would be useful for designing target specific 14-3-3 recognition motif peptides to block interaction of *Pf*14-3-3I with its cognate proteins in the parasite, and develop as potential antimalarial strategy.

## Experimental procedures

Unless stated, all materials were purchased from Sigma-Aldrich, St Louis, MO, USA.

### Culture of P. falciparum

Cryopreserved *P. falciparum* parasites (3D7 laboratory strain) were thawed and cultured according to the protocol as described by W. Trager & JB. Jensen (1976) (51). Briefly, parasite cultures were maintained in O^+^ erythrocytes at 2% hematocrit level, in RPMI 1640 medium (Gibco®, USA) supplemented with 0.5% AlbuMAX^™^ I (Gibco®, USA), 50 mg/L hypoxanthine, 10 mg/L gentamycin (Gibco®, USA) and 2 gm/L sodium bicarbonate. Parasite culture was maintained in ambient hypoxic environment (5% O_2_ and 5% CO_2_; balanced with N_2_). Prior to setting up an experiment, parasite culture was tightly synchronized at ring stage by lysing erythrocytes parasitized with mature parasites, with 5% sorbitol, for two successive intra-erythrocytic proliferative cycles of the parasite.

### Comparative sequence analysis and domain architecture of Pf14-3-3I

Multiple Sequence Alignment (MSA) was performed on comprehensive ensembles of 14-3-3 protein sequences, contextualizing the patterns of conservation and correlation observed in *Pf*14-3-3I protein sequence in light of the structurally and functionally well-characterized orthologs in humans. Amino acid sequences of 14-3-3 isoforms from *Homo sapiens* (seven isoforms: 14-3-3 epsilon, P62258; beta/alpha, P31946; zeta/delta, P63104; theta, P27348; gamma, P61981; eta, Q04917 and sigma, P31947) and *P. falciparum* strain 3D7 (two isoforms as annotated by database curators: *Pf*14-3-3I, PF3D7_0818200 and *Pf*14-3-3II, PF3D7_1362100) were retrieved from Universal Protein Resource (UniProt; https://www.uniprot.org/) and PlasmoDB Plasmodium Genomic Resource (https://plasmodb.org/plasmo/) databases, respectively (52, 53). MSA was done by using MultAlin tool, which creates a multiple sequence alignment from a group of related sequences by using progressive pairwise alignments. (http://multalin.toulouse.inra.fr/multalin/) (54). Based on literature review, and comparative sequence analysis with the *Hs*14-3-3 isoforms, probable amino acid residues of *Pf*14-3-3I involved in dimerization and phosphopeptide (target) binding were utilized to draw overall *Pf*14-3-3I architecture (26, 30, 49, 55, 56). This was done by using Illustrator for Biological Sequences (IBS 1.0.3), a tool for visualizing biological sequences (http://ibs.biocuckoo.org/) (57).

### Phylogenetic analysis and 14-3-3 binding consensus motifs

Amino acid sequences of *Pf*14-3-3 isoforms were compared with their orthologs present across all three kingdoms of life: animalia, plantae & fungi, and a phylogenetic relationship were established. To obtain full-length 14-3-3 protein sequences, BLASTp (https://blast.ncbi.nlm.nih.gov/Blast.cgi) analysis was performed by using amino acid sequence of *Pf*14-3-3I protein as *query sequence*, and proteomes of animals (taxid:33208), plants (taxid:3193) and fungi (taxid:4751) as *search sets*. Publicly available databases such as Uniprot (https://www.uniprot.org/) and The Arabidopsis Information Resource (https://www.arabidopsis.org/) were also utilized to retrieve 14-3-3 protein sequences from *Arabidopsis thaliana* and *Homo sapiens* (52, 58). Literature survey was also done to identify 14-3-3 proteins from *Saccharomyces cerevisiae* and certain plant species including *Solanum lycopersicum, Populus trichocarpa, Oryza sativa, Glycine max* & *Medicago trucatula* (59–61). 14-3-3 protein sequences, thus retrieved from the BLASTp search, databases and literature survey (Supporting file S1) were further categorised into two groups: *epsilon* and *non-epsilon*. Unrooted phylogenetic relationship of *Pf*14-3-3 isoforms with their orthologs was established by using MEGA6, a user-friendly software suite for analysing DNA and protein sequence data from species and populations (http://www.megasoftware.net/) (62).

To identify experimentally validated 14-3-3 interacting partners from prokaryotes & eukaryotes, and collate details of phosphoSer/Thr sites on the target proteins that have been reported to bind directly to 14-3-3 proteins, literature survey followed by mining of publically available databases was done. The Eukaryotic Linear Motif resource (ELM), a database of experimentally validated eukaryotic linear motifs (http://elm.eu.org/), and ANnotation and Integrated Analysis of the 14-3-3 interactome database (ANIA), which integrates multiple data sets on 14-3-3 binding phosphoproteins (https://ania-1433.lifesci.dundee.ac.uk/prediction/webserver/index.py) were utilized for this purpose (63, 64). The repertoire of experimentally determined gold-standard 14-3-3 binding phosphoSer/Thr sites was further extended from the literature survey to give a list of 323 mode I sites from 243 target proteins, 81 mode II sites from 77 target proteins and 9 mode III sites from 9 target proteins (Supporting table S1) (65–67). Amino acid sequences of the 14-3-3 target phosphopeptides were used to *update* optimal consensus 14-3-3 binding motifs by using WebLogo 3, a web based application designed to generate sequence logos (http://weblogo.berkeley.edu/) (68). Protein kinases (CDPK1, PKG, PKA_R_ and PKA_C_) were filtered out as putative binding partners of *Pf*14-3-3I by combining 14-3-3 binding *sequence motifs search* and publicly available collection of annotated phosphoSer/Thr sites gathered from global phospho-proteomic datasets of peptides enriched from schizont stage of *P. falciparum* (69–73). Overall domain architectures of the protein kinases were drawn by using Illustrator for Biological Sequences (IBS 1.0.3) (57).

### cDNA preparation, molecular cloning, over-expression & purification of recombinant 14-3-3I and CDPK1 proteins

cDNA from *P. falciparum* schizonts was prepared by following a previously described protocol, in the absence and presence of Reverse Transcriptase (RT) (74). Integrity of cDNA, thus obtained, was checked by amplification of transcripts encoding for 18S rRNA with the following primer sets: *18S*_Fwd, 5’-CCGCCCGTCGCTCCTACCG-3’ and *18S*_Rev, 5’-CCTTGTTACGACTTCTCCTTCC-3’. 18S rRNA was also amplified from genomic DNA (gDNA) of the parasite, as positive control. Presence of *Pf14-3-3I* transcripts in the cDNA was confirmed by amplification with the following primer sets: *Pf14-3-3I*_Fwd, 5’-TTGTACTTATCGTTCCAAA-3’ and *Pf14-3-3I*_Rev, 5’-TTTCTGAAGTGTTTGGAAT-3’. The experiment was done twice. DNA fragment encoding full-length *Pf*14-3-3I protein was amplified with the following primer sets: *Pf14-3-3I*_Fwd, 5′-TGCGGGATCCATGGCAACATCTGAAGAATTAAAACA-3′ and *Pf14-3-3I*_Rev, 5′-ATTTGTCGACTCATTCTAATCCTTCGTCTTTTGATT-3′ by using Phusion high-fidelity DNA polymerase (Thermo Scientific) and cDNA prepared from the parasites as template. Amplified DNA fragment encoding *Pf*14-3-3I was cloned between BamHI and SalI sites in pGEX-4T-1 vector (with a cleavable Glutathione S-Transferase tag at the N-terminus). Recombinant plasmid was transformed into *Escherichia coli* DH5α competent cells, and positive clones were confirmed by restriction digestion of recombinant plasmid and Sanger sequencing of insert DNA fragment.

For over-expression of GST-14-3-3I fusion protein, the recombinant plasmid containing *14-3-3I* gene was transformed into *E. coli* strain Rosetta competent cells. Over-expression of the protein was induced with 0.5 mM IsoPropyl β-D-1-ThioGalactopyranoside (IPTG) at 0.4 - 0.5 Optical Density (OD; at 600 nm) of bacterial secondary culture, for 8 hours at 25°C. Cells were harvested by centrifugation at 4,000g in Sorvall® Evolution™ RC Centrifuge (Thermo Fisher Scientific™) at 4 °C for 15 minutes. Bacterial pellet, thus obtained, was resuspended in cell lysis buffer (buffer A) containing 10 mM HEPES-NaOH, pH 7.4; 1 mM Ethylene Diamine Tetraacetic Acid (EDTA); 150 mM NaCl; 25μg/ml lysozyme; 3 mM β-MercaptoEthanol (βME), Protease Inhibitor Cocktail (PIC, Roche) and 1mM PhenylMethylSulfonyl Fluoride (PMSF), followed by sonication for 15 minutes with successive pulses of 6 seconds ON and 10 seconds OFF. Cell lysate was centrifuged at 15,000 g at 4 °C for 1hr. Supernatant was loaded onto Glutathione Sepharose High Performance resin packed column, GSTrap™ HP 1-mL (GE Healthcare) pre-equilibrated with buffer B (10 mM HEPES-NaOH, pH 7.4; 1 mM EDTA and 150 mM NaCl), for overnight at 4 °C. Column was washed with 10 fold column volume of buffer B and recombinant 14-3-3I protein was eluted with different concentrations (10, 20 & 50 mM) of reduced glutathione, prepared in buffer B. Elution fractions, thus obtained, from affinity purification were pooled and buffer exchanged using Amicon™ Ultra-15 Centrifugal Filter Unit (10 kDa cutoff; Merck™) in 10 mM HEPES-NaOH buffer, pH 7.4 and 150 mM NaCl.

To raise specific antibodies against r14-3-3I protein, Balb/c mice were immunized with 50 μg of r14-3-3I protein in 0.9% saline. The formulation was made by thoroughly mixing equal volumes of Freund’s complete adjuvant and saline containing r14-3-3I protein. For booster doses, formulations were made with Freund’s incomplete adjuvant. Once the anti-serum was raised, native *Pf*14-3-3I protein was detected in schizonts lysate by immunoblotting. The experiment was done twice.

Cloning of *cdpk1* gene in pET-28a(+) vector (with 6x-His tags at N- and C-termini) was done as described earlier (74). For over-expression of 6x-His-CDPK1 fusion protein, the recombinant plasmid harboring *cdpk1* gene was transformed into *E. coli* strain BLR (λDE3) competent cells. Over-expression of the protein was induced with 1 mM IPTG at 0.9 OD of bacterial secondary culture, for 4 hours at 30°C. Protein purification was done in a similar manner as the purification of r14-3-3I, except for the following changes in buffers’ compositions. Buffer A: 10 mM Tris, pH 8.0; 1 mM EDTA; 100 mM NaCl; 25μg/ml lysozyme; 3 mM βME and PIC. Buffer B: 10 mM imidazole; 10 mM Tris, pH 8.0; 100 mM NaCl and 3 mM βME. Supernatant obtained after centrifugation of bacterial lysate was loaded onto High-Performance Immobilized Metal Affinity Chromatography (IMAC) column, HisTrap™ HP 1-mL (GE Healthcare) and recombinant CDPK1 protein was eluted with different concentrations of imidazole, ranging from 20-500 mM, prepared in buffer B. Elution fractions, thus obtained, from metal-affinity purification were pooled and buffer exchanged, as mentioned above.

### Immuno-Fluorescence Assay (IFA) to detect expression and co-localization of 14-3-3I with CDPK1

IFAs were performed on synchronized *P. falciparum* 3D7 culture to check for expression of 14-3-3I and its co-localization with CDPK1 in mature stages of the parasite, as described earlier (75). Briefly, mature schizonts (44-46 hpi) & merozoites were isolated, smeared on glass slides, dried and fixed with pre-chilled methanol for overnight at −20°C. Non-specific binding sites in the parasite were blocked with 3% BSA (in 1X PBS) for 1 hour at room temperature (RT, 293K) and probed with anti-*Pf*14-3-3I mouse serum and anti-*Pf*CDPK1 rabbit serum at dilutions of 1:50, for 1 hour at RT. Slides were washed twice with PBS containing 0.05% Tween-20 (PBST) followed by washing once with PBS, and probed with Alexa-Fluor 488 conjugated anti-mouse IgG (Molecular Probes, USA) and Alexa-Fluor 594 conjugated anti-rabbit IgG, both at dilutions of 1:500 for 1 hour at RT. All antibodies were diluted in 1% BSA, prepared in 1X PBS. The slides were washed and mounted with ProLong Gold antifade reagent with DAPI (4’,6-diamidino-2-phenylindole) (Invitrogen) and images were acquired using Nikon A1-R confocal microscope using the NIS Elements software. For clear visual assessments of (co)localization, images were processed by Imaris 7.0.0 (Bitplane Scientific), which provides functionality for visualization, segmentation and interpretation of 3D microscopy data sets. The experiment was done thrice.

### Homology modeling of Pf14-3-3I and PfCDPK1

Three-dimensional structure of a protein can provide us with precise information about its single, most stable conformation, as dictated by its sequence. Comparative or homology modeling, one of the most common structure prediction methods in structural genomics and proteomics, was employed to model 3D structures of 14-3-3I_dimer_ and CDPK1 from *P. falciparum* strain 3D7. To accomplish this feat, amino acid sequences of 14-3-3I (PF3D7_0818200) and CDPK1 (PF3D7_0217500) were retrieved from PlasmoDB database (53). To search for a suitable template for homology modeling, BLASTp (https://blast.ncbi.nlm.nih.gov/Blast.cgi) search was performed by using amino acid sequences of 14-3-3I and CDPK1 as *query sequences*, against Protein Data Bank (PDB) database (http://www.rcsb.org/) (76, 77). Amino acid sequence identity between 14-3-3 orthologs from *P. falciparum* strain 3D7 (14-3-3I) and *Homo sapiens* (14-3-3 Epsilon) was found to be 63.52%, thus rendering X-Ray diffraction based structural model for *Hs*14-3-3 epsilon (PDB ID: 3UAL; resolution: 1.8 Å) as a suitable template to model 3D structure of *Pf*14-3-3I_dimer_ (78). For *Pf*CDPK1, X-ray diffraction based structural model of CDPK1 protein from *P. berghei* strain ANKA (PDB ID: 3Q5I; sequence similarity: 93%) was used as a suitable template (79). Homology modeling was done by using Modeller v9.17 software (https://salilab.org/modeller/), a program designed for comparative protein structure modeling by satisfaction of spatial restraints (80). The best structural model with the most negative DOPE score was selected and rendered with PyMOL Molecular Graphics System, v2.1 by Schrödinger, LLC (http://pymol.org/2/) (81).

Generated structural models were further subjected to structural refinement by using ModRefiner (https://zhanglab.ccmb.med.umich.edu/ModRefiner/) which is an algorithm-based approach for atomic-level, high-resolution protein structure refinement (82). Reliability of the refined structural models of *Pf*14-3-3I_dimer_ and *Pf*CDPK1 was assessed by examining backbone dihedral (torsion) angles: phi (Ø) and psi (Ψ) of the amino acid residues lying in the energetically *favourable* regions of Ramachandran space (83). This was done by using online available tool, PROCHECK v.3.5 (https://www.ebi.ac.uk/thornton-srv/software/PROCHECK/) (84). Percentage quality measurement of the protein structures was evaluated by utilizing four sorts of occupancies called ‘core’, ‘additional allowed’, ‘generously allowed’ and ‘disallowed’ regions. The refined 3D structural models of *Pf*14-3-3I_dimer_ and *Pf*CDPK1, thus generated, were subsequently used for *Pf*14-3-3I_dimer_/*Pf*CDPK1 PPI & *Pf*14-3-3I_dimer_/peptides interaction and molecular simulation studies.

### Molecular Docking

Molecular docking of 14-3-3I_dimer_ against the structurally modeled phosphopeptides (peptide 1: ARSH*pS*YPA and peptide 2: RLYH*pS*LPA; *pS*: phosphorylated serine) was performed by using AutoDock 4.2.6 tool (http://autodock.scripps.edu/) (85). For the receptor 14-3-3I_dimer_ protein, we assigned polar Hydrogen and Kollman charges, whereas Marsilli-Gasteiger partial charges were assigned for the peptides. We have not used any additional restraint on torsion angles of the receptor protein during the docking process. The Lamarckian Genetic Algorithm (LGA) was used for performing the energy evaluations with a default set of docking parameters. All the docked complexes were evaluated based on their lowest binding energy (kcal/mol).

CDPK1 was docked against 14-3-3I_dimer_ receptor by using High Ambiguity Driven biomolecular DOCKing (HADDOCK) webserver, an information-driven flexible docking approach for the modeling of biomolecular complexes (86). We preferentially selected those Serine and Threonine residues that are previously reported to be autophosphorylated by CDPK1. To phosphorylate them *in silico*, we adopted the PyTMs plugin in PyMOL Molecular Graphics System (87, 88). During the whole docking process, the phosphorylated Serine and Threonine residues, present at the interaction surface of CDPK1 and 14-3-3I_dimer_ receptor, were considered as *active* residues and the amino acid residues surrounding those as *passive* residues.

### Molecular Dynamics (MD) simulation Preparation and Parameterization

14-3-3I_dimer_/CDPK1 & 14-3-3I_dimer_/peptides complex systems were parameterized by using terminal interface, *tLeap* of Assisted Model Building with Energy Refinement tool, AMBER 14 (89). The system was neutralized by using the counter-charged ions (Na^+^ or Cl^-^) followed by solvation using explicit TIP3P water model spanning for 12 Å around the complex systems. The Forcefield *ff03*.*r1* was used for the proteins & peptides, and the phosphorylated Serine and Threonine residues were handled by using the *ffptm* force filed (90, 91).

### Simulation

14-3-3I_dimer_/CDPK1 & 14-3-3I_dimer_/peptides complex systems were equilibrated by carrying out a short minimization (1000 steps, 500 cycles of steepest descent and 500 cycles of conjugate gradient minimization) runs followed by 50ps of heating, 50ps of density equilibration and 500ps of constant pressure equilibration, performed at 300 K. During the equilibration process, the bonds involving Hydrogen-atoms were handled by SHAKE algorithm (92). The simulations were carried out at 1fs time step with Langevin thermostat, under the “periodic boundary conditions” using *sander*, an AMBER 14 module which carries out energy minimization, molecular dynamics, and NMR refinements.

After the equilibration, production runs were performed at 300K, with Langevin thermostat, under a constant pressure. We performed 20ns long simulations for 14-3-3I_dimer_/peptides complexes and 100ns long simulation for 14-3-3I_dimer_/CDPK1 complex, and the coordinates were updated after every 5ps. We used a non-bonded cutoff distance of 8Å, and Particle Mesh Ewald (PME) algorithm handled the long ranged electrostatic interactions (93). The net translational and rotational velocities were cancelled after every 1000 steps. The simulations were carried out at 1fs time step by using *pmemd*, a version of *sander* that is optimized for speed and parallel scaling.

### Analysis and binding energy calculation

Residue specific free energy decomposition was performed by using Molecular Mechanics/Generalized Born and Surface Area (MM/GBSA) continuum solvation method (41). Here, we calculated Gibbs free energy of binding (Δ*G*_bind_) for both 14-3-3I_dimer_/peptides and 14-3-3I_dimer_/CDPK1 interactions, as follows:

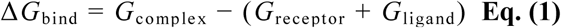

Where, Gibbs free energy is the sum of:

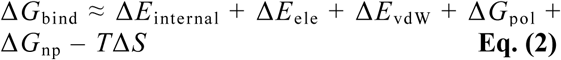

Where, Δ*E*_internal_ includes bond, bend and dihedral energies of the system; Δ*E*_ele_ & Δ*E*_vdW_ denote electrostatic and Van Der Waals interaction energies, respectively; Δ*G*_pol_ & Δ*G*_np_ denote polar and non-polar solvation terms, respectively. The polar solvation term was calculated by Generalized Born (GB) model, whereas the non-polar solvation term was calculated from linear relation to the solvent accessible surface area. *T*Δ*S* is the product of temperature and entropy terms. We used MMPBSA.py.MPI module of AMBER 14 to calculate Δ*G*_bind_ for both 14-3-3I_dimer_/peptides and 14-3-3I_dimer_/CDPK1 interaction systems by using MM/GBSA method (94).

### ELISA to confirm 14-3-3I/CDPK1 interaction

CDPK1 phosphorylation was achieved by setting up a kinase reaction *in vitro* with 100 ng/µl of rCDPK1, in assay buffer [100 mM Tris-Cl, pH 7.4; 2.5 mM DiThioThreitol (DTT); 50 mM MgCl_2_ and 2.5 mM MnCl_2_]. Enzymatic reaction was carried out in a total volume of 200 µl, in the absence & presence of Ca^2+^-ions for conditions requiring minimally & maximally phosphorylated rCDPK1 protein, respectively. 2.5 mM EGTA was added for condition requiring absence of Ca^2+^-ions. Kinase reactions were initiated by adding 2 mM ATP and allowed to take place at 30°C for 1 hr. In all experiments, minimally & maximally phosphorylated CDPK1 protein is represented as rCDPK1 and r*p*CDPK1, respectively.

ELISA was performed to confirm 14-3-3/CDPK1 interaction by following previously described protocol (95). Briefly, purified rCDPK1 protein (200 ng/100 μL/well of rCDPK1 or r*p*CDPK1) was coated onto Poly-L-Lysine coated 96-welled microtitre plate in 0.06 M carbonate-bicarbonate buffer, pH 9.6, for overnight at 4°C. The plate was blocked with 5% skimmed milk solution (in PBS; 200 μL/well) for 1 hour at 37°C and washed thrice with 0.05% Tween in PBS (PBST) for 5 minutes each. For interaction analysis, rCDPK1 coated wells were incubated with different concentrations of r14-3-3I protein: 0.5, 1, 2, 4 and 8 μM (100 μL/well), for 1 hour at 37°C. Following three washes with PBST, anti-GST HRP Conjugated antibody (Sigma Aldrich; diluted 1:5,000 in PBS) was added to each well (100 μL/well) and incubated for 1 hour at 37°C. The peroxidase reaction was developed with o-Phenylenediamine Dihydrochloride (OPD; 1mg/mL) as chromogen and hydrogen peroxide as substrate, both prepared in citrate phosphate buffer, pH 5.0 (100 μL/well). The enzymatic reaction was stopped with 2N H_2_SO_4_ and the optical density was measured spectrophotometrically by taking absorbance at 490 nm using Varioskan™ LUX multimode microplate reader (Thermo Fisher Scientific™). For conditions requiring presence of peptides 1 & 2 as 14-3-3I/CDPK1 interaction inhibitors, ELISA was performed in the presence of different concentrations of the peptides: 1, 5, 10, 25 and 50 μM. The experiment was done twice in triplicates.

### Pull-down assay to confirm 14-3-3I/CDPK1 interaction

To further confirm 14-3-3I/CDPK1 interaction, western blot based GST pull-down assay was performed in which Glutathione Sepharose® 4B beads (GE Healthcare) was coupled with 10 µg of purified r14-3-3I protein in buffer A (20 mM Tris, pH 7.4; 0.5 mM DTT; 10 mM MgCl_2_; 0.5 mM MnCl_2_; 50 mM NaCl; 4% (v/v) glycerol; 0.1 mg/mL BSA and 25 mM CaCl_2_), for 1 hour at RT., followed by washing three times (at 900 rpm for 2 min.) with buffer A. Equivalent amount of purified GST protein was taken as negative control. rCDPK1 protein was auto-phosphorylated in an *in vitro* kinase reaction, as mentioned above. The r14-3-3I beads were incubated with purified rCDPK1 protein (10 µg of rCDPK1 or r*p*CDPK1 per binding reaction), for 1 h at RT. After washing three times with buffer B (buffer A, 300 mM NaCl), the bead-bound protein complexes were boiled in SDS-loading dye and analyzed by SDS–PAGE, followed by immunoblotting with HRP-conjugated anti-His antibody (Sigma-Aldrich) by using SuperSignal™ West Femto Maximum Sensitivity Substrate (Thermo Scientific). The blot was stripped and re-probed with HRP-conjugated anti-GST antibody to check for equal coupling of r14-3-3I (or equivalent amount of GST) with Glutathione Sepharose beads in all binding reactions. For conditions requiring presence of peptides 1 & 2 as 14-3-3I/CDPK1 interaction inhibitors, GST pull-down assay was performed in the presence of 10 μM concentration of the peptides. The blots shown in the figure are representative of two independent experiments.

### Surface Plasmon Resonance analysis of binding affinity between CDPK1 and immobilized 14-3-3I

rCDPK1 protein was auto-phosphorylated in an *in vitro* kinase reaction, as mentioned above. To determine binding strength of CDPK1 and 14-3-3I protein, real-time biomolecular interaction analysis with Surface Plasmon Resonance (SPR) was carried out at physiologically relevant concentrations, by using AutoLab Esprit SPR (at Advanced Instrumentation Research Facility, Jawaharlal Nehru University, New Delhi, India). Kinetic rate constants: K_a_ and K_d_ (association and dissociation rate constants, respectively) as well as affinity constant, K_D_ were measured at RT. SPR analysis was performed by following previously described protocols (96–100). Briefly, 5 μM of recombinant 14-3-3I protein was immobilized on the surface (self-assembled monolayer of 11-Mercapto-Undecanoic Acid, MUA on gold surface) of SPR sensor chip by the mechanism of *covalent amine coupling*. The chemistry involves covalent linkage of esterified carboxyl groups of the sensor surface with ε-amino group of lysine residues of a given protein. Unreacted ester groups on the sensor chip surface were blocked with 100 mM Ethanolamine, pH 8.5. Interaction kinetics was studied by injecting recombinant CDPK1 or *p*CDPK1 protein over the 14-3-3I immobilized chip surface at different concentrations: 100, 250, 500, 750 and 1000 nM, at a steady rate of 20 μL/min, with association and dissociation time of 300s and 150s, respectively. HEPES-NaOH buffer was used both as immobilization and binding solutions. Surface of the sensor chip was regenerated with 50 mM NaOH solution. The experiment was done twice.

Data were fit to the two-state conformational change model by using AutoLab SPR Kinetic Evaluation software provided with the instrument. At least three independent experiments were performed. K_D_ value was calculated by using the Integrated Rate Law (IRL) equation:

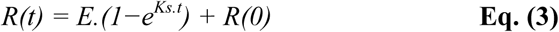

where, E is the maximal extent of change in response at a certain concentration and is equal to k_a_.C.R_max_/(k_a_.C + k_d_); K_s_ is equal to (k_a_.C+ k_d_); and R(0) is the response unit at *t=0*. E and K_s_ were evaluated at each concentration by minimizing residual sum of squares between observed data and the model equation using solver in MS Excel. Evaluation of K_s_ at four different concentrations of the compounds was done to calculate K_D_ according to the following equation:

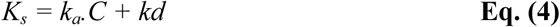

where, K_s_ is a concentration dependent parameter which is dependent upon k_a_ and k_d_. Ratio of intercept (k_d_) and slope (k_a_) of the above line was ascertained to be the dissociation constant or K_D_.

### Isothermal Titration Calorimetric analysis

To calculate kinetic parameters such as binding affinity constant (K_a_) for interaction of 14-3-3I with consensus 14-3-3 binding peptides 1 and 2, Isothermal Titration Calorimetry (ITC) experiments were performed by using MicroCal iTC200 (Malvern Instruments Ltd, UK; at School of Physical Sciences, Jawaharlal Nehru University, New Delhi, India). For this purpose, recombinant 14-3-3I protein was dialyzed extensively against HEPES-NaCl buffer (10 mM HEPES-NaOH, pH 7.4 and 150 mM NaCl) using Amicon^™^ Ultra-15 Centrifugal Filter Unit (10 kDa cutoff) before its subsequent use in ITC. Buffer was degassed by vacuum for 10 minutes prior to use. Dilutions of r14-3-3I and peptides were prepared in HEPES-NaCl buffer to ignore contribution from buffer-buffer interaction. ITC analysis was done at RT, by following previously described protocols (99, 101, 102). Briefly, syringe was loaded with 40 μL of the ligand (i.e., peptide 1 or 2) at concentration of 40 μM (peptide 1) & 100 μM (peptide 2), and sample cell was filled with 280 μL of r14-3-3I protein at 10 μM concentration. Volume of first injection of the ligand was set to 0.4 μL, with an initial delay of 60 sec. It was followed by 19 successive injections each of 2μL, with an interval of 150 seconds between each injection to allow the signals to reach baseline. Background titration profiles were obtained by injecting ligand into the buffer solution, under identical experimental conditions. Heat of dilution of the ligand, thus obtained, was subtracted to determine net enthalpy change for 14-3-3I/(peptide 1 or 2) interaction. The experiment was done twice. Amount of heat produced per injection (corrected data) was analyzed by integration of area under individual peaks by MicroCal ORIGIN 7 software provided by the instrument manufacturer. Experimental data was presented as the amount of heat produced per second (μcal/sec; corrected for heat of dilution of the ligand) following each injection of the ligand into the protein solution, as a function of time (minutes). Interaction data was represented as the best fit of the non-linear experimental data to the single-site binding model yielding molar binding stoichiometry (N), binding constant (K_a_), enthalpy change (Δ*H*) and entropy change (Δ*S*). Using K_a_, Δ*H* and Δ*S*, the Gibbs free energy change (Δ*G*) was calculated by using the following equation:

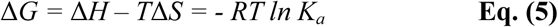

### Evaluation of growth inhibitory effect on malaria parasites

Phosphopeptides 1 & 2 were subjected to parasite growth inhibition *in vitro*, as described previously (103). Briefly, mature (punctated) schizonts at 1% initial parasitaemia were treated with 12.5 μM concentration of the peptides, for 24 hours at 37°C. Untreated parasites were taken as control. After progression of the parasites to ring stage, erythrocytes were washed with 1X PBS and stained with Ethidium Bromide (EtBr, 10 μM) for 20 minutes at RT, in dark. Following two washes with 1X PBS, cells were analyzed by flow cytometry on BD LSRFortessa™ cell analyzer using FlowJo v10 software. Fluorescence signal (FL-2) was detected with the 590 nm band pass filter by using an excitation laser of 488 nm collecting 100,000 cells per sample. Following acquisition, parasitaemia levels were estimated by determining the proportion of FL-2-positive cells using Cell Quest. The experiment was done twice in triplicates. Percent parasite growth inhibition was calculated as follows.

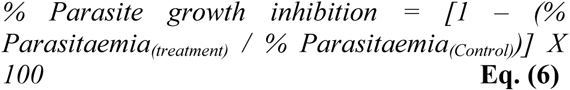

## Statistical analysis

In the bar graphs, data is expressed as Mean ± Standard Deviation (SD) of three independent experiments, done in duplicates. Statistical analysis was done by using OriginPro Evaluation 2018b Graphing and Analysis software. Unless indicated, the differences were considered to be statistically significant at P < 0.05.

## Acknowledgements

Authors acknowledge AutoLab Esprit SPR facility of Advanced Instrumentation Research Facility (AIRF), Jawaharlal Nehru University (JNU), New Delhi, India and Central Instrumentation Facility (CIF) of SCMM, JNU for flow cytometry. The lab facility of Shiv Nadar University is also acknowledged.

## FOOTNOTES

This work has been funded by DST-EMR from the Department of Science and Technology (DST), Ministry of Science and Technology, Government of India. Shailja Singh is a recipient of the Innovative Young Biotechnologist Award (IYBA) from Department of Biotechnology (DBT). RJ is supported from University Grants Commission-Junior Research Fellowship (UGC-JRF).

