## Supporting information for "Interaction of 14-3-3I and CDPK1 mediates the growth of human malaria parasite"

***List of the material included***

**Supporting figure S1:** Homology model of *Pf*14-3-3$I_{dimer}$ and its quality assessment.

**Supporting figure S2:** Evolutionary relationship of *Pf*14-3-3 isoforms with its orthologs from kingdom plantae, animalia and fungi.

**Supporting figure S3:** Updated 14-3-3 binding consensus motifs.

**Supporting table S1:** Updated repertoire of experimentally validated 14-3-3 binding phosphoSer/Thr sites on target proteins from prokaryotes and eukaryotes, as identified by literature survey and mining of publically available databases.

**Movie S1:** Movie showing the interactions between the *Pf*14-3-$3_{dimer}$ protein (blue) and *p*CDPK1 (grey). The phosphoSer and phosphoThr residues of *p*CDPK1 are shown in red and yellow, respectively.

**Movie S2:** Movie showing the interactions between the *Pf*14-3-$3_{dimer}$ protein (blue) and peptide 1 (yellow). The phosphoSer of peptide 1 is shown in red.

**Movie S3:** Movie showing the interactions between the *Pf*14-3-$3_{dimer}$ protein (blue) and peptide 2 (yellow). The phosphoSer of peptide 2 is shown in red.

**Supporting file 1:** 14-3-3 protein sequences retrieved from BLASTp search, databases' mining and literature survey.

### Figure S1

**A.**

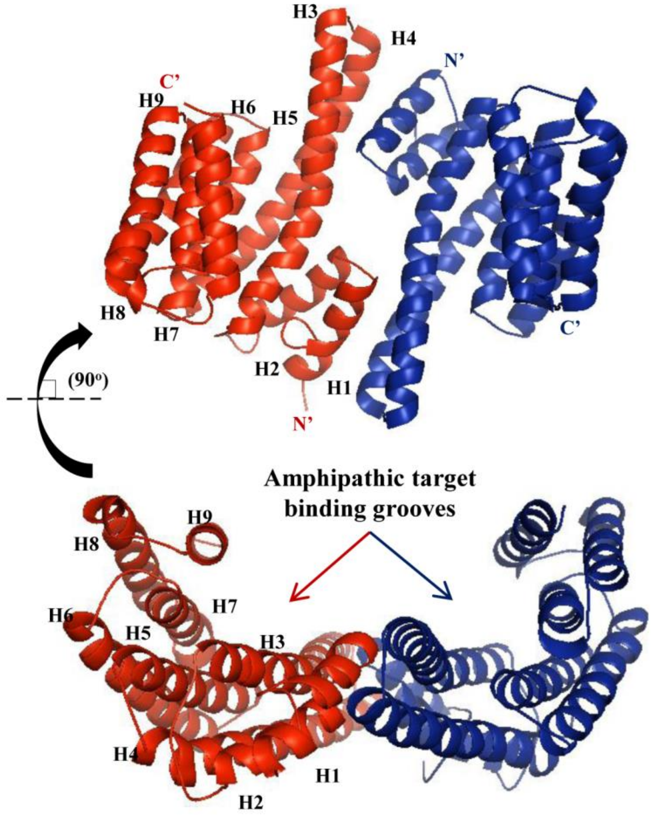
H3
H4
N'
C'
H9
H6
H5
H8
H7
H2
H1
N'
C'
(90°)
Amphipathic target
binding grooves
H9
H8
H7
H6
H5
H3
H4
H1
H2

**B.**

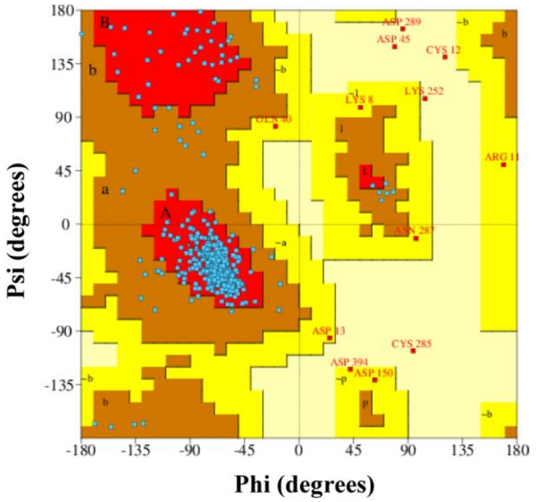
Psi (degrees)
Phi (degrees)
180
135
90
45
0
-45
-90
-135
-180
-135
-90
-45
0
45
90
135
180
ASP 289
ASP 45
CYS 12
LYS 8
LYS 252
ARG 11
ASP 13
CYS 285
ASP 394
ASP 150

**Supporting figure S1: Homology model of *Pf*14-3-3I$_{\mathbf{dimer}}$ and its quality assessment. A)** *Comparative structural model of 14-3-3I$_{dimer}$*. X-Ray diffraction based structural model of *Hs*14-3-3 epsilon (PDB ID: 3UAL) was used as template to model 3D-structure of 14-3-3I$_{dimer}$ by using Modeller v9.17. Generated model was subjected to structural refinement by using ModRefiner. Overall RMSD value of the C-alpha atomic coordinates, upon optimal rigid-body superimposition of *Hs*14-3-3 epsilon and *Pf*14-3-3I$_{dimer}$ was found to be 0.63 Å, suggesting a reliable structural model of *Pf*14-3-3I$_{dimer}$. Structural folds depicted a clamp like homo-dimer where each monomer harbored amphipathic groove for binding to (target) phosphorylated sequence motifs. Also, both monomers of *Pf*14-3-3I were found to be oriented in opposite direction. Helical regions in one of the monomers are marked from H1 to H9. **B)** *Stereochemical assessment of the generated homology model.* Assessment of backbone dihedral (torsion) angles: phi (Ø) and psi (Ψ) of the amino acid residues displayed 88.7% of the residues lying in the most favored ("core") regions, with 8.8%, 1.5%, and 1.1% residues in "additional allowed", "generously allowed" and "disallowed regions" of Ramachandran plot, respectively. This was done by using PROCHECK v.3.5.

### Figure S2

**A.**

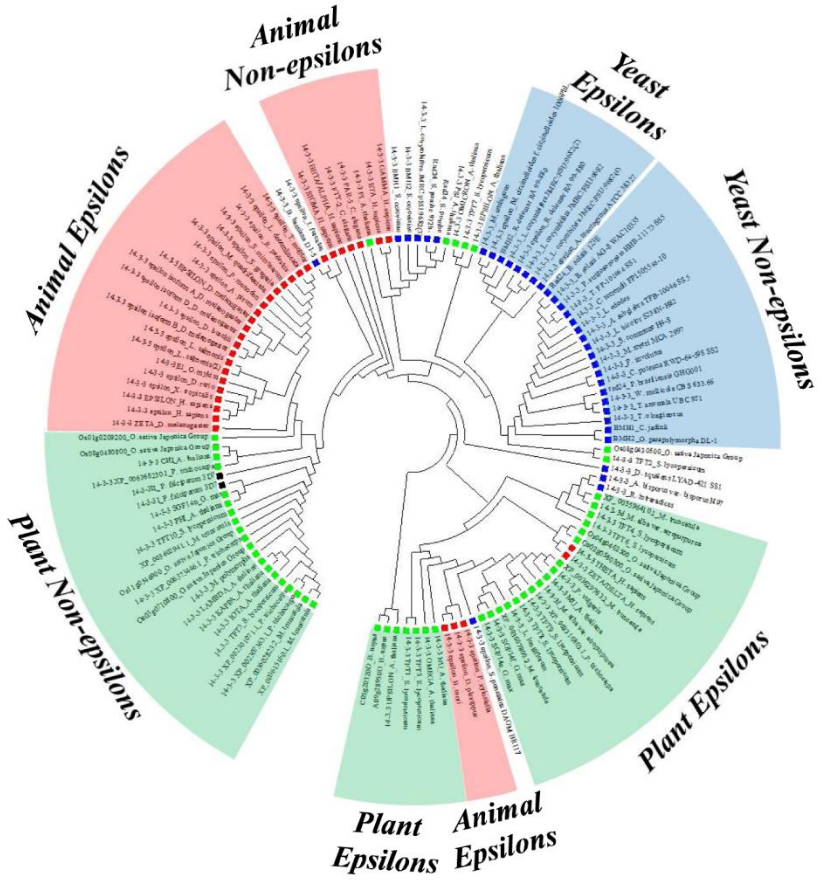


**B.** **Plant NON-EPSILON Group**

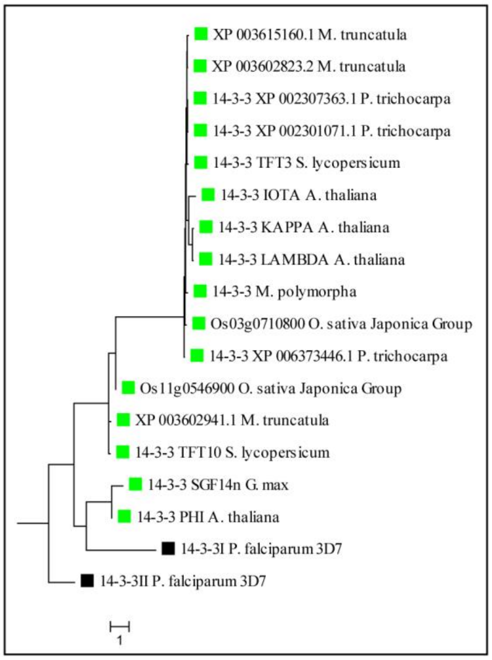

**Supporting figure S2: Evolutionary relationship of *Pf*14-3-3 isoforms with its orthologs from kingdom plantae, animalia and fungi. A)** *Unrooted phylogenetic relationship.* Phylogenetic analysis showed that *Pf*14-3-3 isoforms have followed convergent evolutionary pathway with 14-3-3 proteins from plant "non-epsilon group". Analysis was done by using MEGA6. Branches with green, red and blue squares belong to kingdom plantae, animalia and fungi, respectively. **B)** Detailed evolutionary relationship of *Pf*14-3-3 isoforms with its orthologs from plant "non-epsilon group" is shown.

### Figure S3

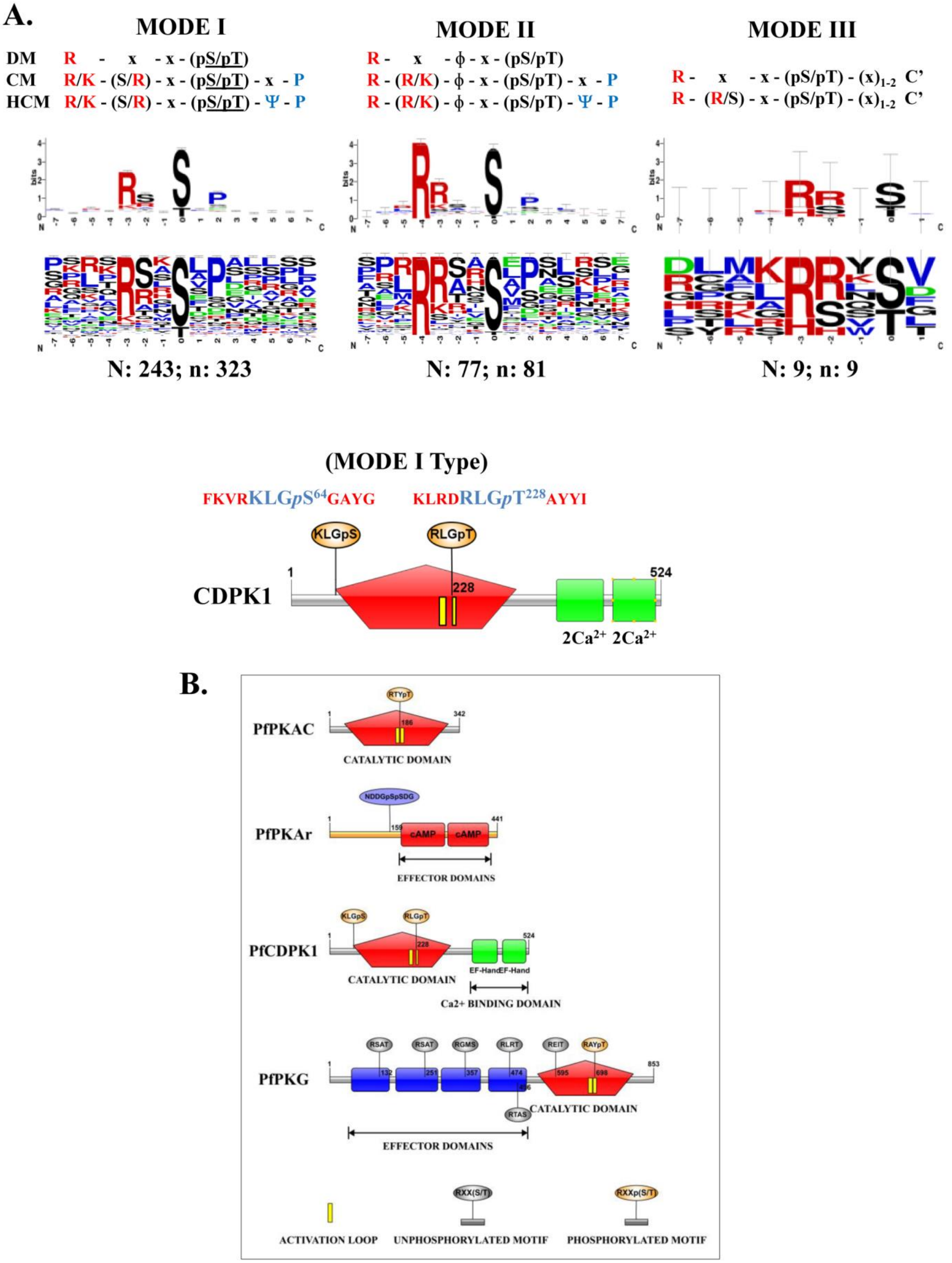
A.
MODE I
DM R - x - x - (pS/pT)
CM R/K - (S/R) - x - (pS/pT) - x - P
HCM R/K - (S/R) - x - (pS/pT) - Ψ - P
MODE II
R - x - ϕ - x - (pS/pT)
R - (R/K) - ϕ - x - (pS/pT) - x - P
R - (R/K) - ϕ - x - (pS/pT) - Ψ - P
MODE III
R - x - x - (pS/pT) - (x)1-2 C'
R - (R/S) - x - (pS/pT) - (x)1-2 C'
N: 243; n: 323
N: 77; n: 81
N: 9; n: 9
(MODE I Type)
FKVRKLGpS64GAYG
KLRDRLGpT228AYYI
KLGpS
RLGpT
CDPK1
1
228
524
2Ca2+ 2Ca2+
B.
PfPKAC
RTYpT
186
342
CATALYTIC DOMAIN
PfPKAr
NDDGpSpSDG
159
441
cAMP
EFFECTOR DOMAINS
PfCDPK1
KLGpS
RLGpT
228
524
EF-Hand EF-Hand
CATALYTIC DOMAIN
Ca2+ BINDING DOMAIN
PfPKG
RSAT
RSAT
RGMS
RLRT
REIT
RAYpT
132
251
357
474
496
595
698
853
RTAS
CATALYTIC DOMAIN
EFFECTOR DOMAINS
RXX(S/T)
RXXp(S/T)
ACTIVATION LOOP
UNPHOSPHORYLATED MOTIF
PHOSPHORYLATED MOTIF

**Supporting figure S3: Updated 14-3-3 binding consensus motifs. A)** Experimentally validated 14-3-3 binding phosphoSer/Thr sites were identified from prokaryotes & eukaryotes by literature survey and mining of publically available databases. Finally, an updated repertoire of 323 mode I sites from 243 target proteins, 81 mode II sites from 77 target proteins and 9 mode III sites from 9 target proteins was obtained. Amino acid sequences of these phosphopeptides were used to update 14-3-3 binding consensus motifs by using WebLogo3. Amino acid sequences of probable 14-3-3 binding phosphopeptides of CDPK1 are shown. **B)** Protein kinases: CDPK1, PKG, $PKA_R$ and $PKA_C$ were filtered out as putative binding partners of *Pf*14-3-3I by combining the updated 14-3-3 binding consensus motifs and publicly available datasets of phosphopeptides from schizonts. Overall domain architectures of the kinases were drawn by using Illustrator for Biological Sequences (IBS 1.0.3). DM: Degenerate Motif; CM: Consensus Motif; HCM: Highly Consensus Motif.
