## Supporting file S1 for "Interaction of 14-3-3I and CDPK1 mediates the growth of human malaria parasite"

**Protein sequences used for phylogenetic analysis of 14-3-3 family proteins from different species covering kingdoms animalia, plantae and fungi**

#### 1. *Plasmodium falciparum* 14-3-3 protein isoforms
**Source: PlasmoDB Database**

Aurrecoechea, C., Brestelli, J., Brunk, B. P., Dommer, J., Fischer, S., Gajria, B., Gao, X., Gingle, A., Grant, G., Harb, O. S., Heiges, M., Innamorato, F., Iodice, J., Kissinger, J. C., Kraemer, E., Li, W., Miller, J. A., Nayak, V., Pennington, C., Pinney, D. F., Roos, D. S., Ross, C., Stoeckert, C. J., Treatman, C., and Wang, H. (2009) PlasmoDB: A functional genomic database for malaria parasites. *Nucleic Acids Res.* **37**, D539–D543

**14-3-3I**
>gi|296005038|ref|XP_002808856.1| 14-3-3 protein, putative [Plasmodium falciparum 3D7]
MATSEELKQLRCDCTYRSKLAEQAERYDEMADAMRTLVEQCVNNDKDELTVEERNLLSVAYKNAVGARRA
SWRIISSVEQKEMSKANVHNKNVAATYRKKVEEELNNICQDILNLLTKKLIPNTSESESKVFYYKMKGDY
YRYISEFSCDEGKKEASNCAQEAYQKATDIAENELPSTHPIRLGLALNYSVFFYEILNQPHQACEMAKRA
FDDAITEFDNVSEDSYKDSTLIMQLLRDNLTLWTSDLQGDQTEEKSKDEGLE

**14-3-3II**
>gi|296005562|ref|XP_002809097.1| 14-3-3 protein, putative [Plasmodium falciparum 3D7]
MNQYIDNDISVSNKEELIYYIKILNHLGSYDETVTLIKSVNVENYNFNYTESLSVGFAFKNALNVKRKEK
TILENIITNEKSSEREKICAELLKSKLNTDIRSIEKNTYAVLKNKCISRTTDDKILMLYWHILGDMSRYC
ADTFHGTDKEKMHEKSMKSYSYALHYANKMKIPPSSPKMLELLVSWTVLHKDMNKDINYSIELAAEAFRN
AIQNMHLLENDDECSKTIRILGILRDNINKWCEISGRKNVNALFEINGENLDKYKDIMNSTHT

#### 2. *Arabidopsis thaliana* 14-3-3 protein gene family
**Source: The Arabidopsis Information Resource**

Berardini, T. Z., Reiser, L., Li, D., Mezheritsky, Y., Muller, R., Strait, E., and Huala, E. (2015) The arabidopsis information resource: Making and mining the "gold standard" annotated reference plant genome. *Genesis*. **53**, 474–85

**CHI**
>gi|1256534|gb|AAA96323.1| GF14 chi chain [Arabidopsis thaliana]
MATPGASSARDEFVYMAKLAEQAERYEEMVEFMEKVAKAVDKDELTVEERNLLSVAYKNVIGARRASWRI
ISSIEQKEESRGNDDHVSLIRDYRSKIETELSDICDGILKLLDTILVPAAASGDSKVFYLKMKGDYHRYL
AEFKSGQERKDAAEHTLTAYKAAQDIANSELAPTHPIRLGLALNFSVFYYEILNSPDRACNLAKQAFDEA
IAELDTLGEESYKDSTLIMQLLRDNLTLWASDMQDDVADDIKEAAPAAAKPADEQQS

**OMEGA**
>gi|553040|gb|AAA32798.1| GF14 [Arabidopsis thaliana]
MASGREEFVYMAKLAEQAERYEEMVEFMEKVSAAVDGDELTVEERNLLSVAYKNVIGARRASWRIISSIE
QKEESRGNDDHVTAIREYRSKIETELSGICDGILKLLDSRLIPAAASGDSKVFYLKMKGDYHRYLAEFKT
GQERKDAAEHTLAAYKSAQDIANAELAPTHPIRLGLALNFSVFYYEILNSPDRACNLAKQAFDEAIAELD
TLGEESYKDSTLIMQLLRDNLTLWTSDMQDDAADEIKEAAAPKPTEEQQ

**PSI**
>gi|166717|gb|AAA32799.1| GF14 psi chain [Arabidopsis thaliana]
MSTREENVYMAKLAEQAERYEEMVEFMEKVAKTVDVEELSVEERNLLSVAYKNVIGARRASWRIISSIEQ
KEESKGNEDHVAIIKDYRGEIESELSKICDGILNVLEAHLIPSASPAESKVFYLKMKGDYHRYLAEFKAG
AERKEAAESTLVAYKSASDIATAELAPTHPIRLGLALNFSVFYYEILNSPDRACSLAKQAFDDAIAELDT

LGEESYKDSTLIMQLLRDNLTLWTSDMTDEAGDEIKEASKPDGAE

**PHI**

>gi|1493805|gb|AAB06231.1| GF14 protein phi chain [Arabidopsis thaliana]
MAAPPASSSAREEFVYLAKLAEQAERYEEMVEFMEKVAEAVDKDELTVEERNLLSVAYKNVIGARRASWR
IISSIEQKEESRGNDDHVTTIRDYRSKIESELSKICDGILKLLDTRLVPASANGDSKVFYLKMKGDYHRY
LAEFKTGQERKDAAEHTLTAYKAAQDIANAELAPTHPIRLGLALNFSVFYYEILNSPDRACNLAKQAFDE
AIAELDTLGEESYKDSTLIMQLLRDNLTLWTSDMQDESPEEIKEAAAPKPAEEQKEI

**UPSILON**

>gi|1508785|gb|AAB06585.1| GF14 upsilon chain [Arabidopsis thaliana]
MSSDSSREENVYLAKLAEQAERYEEMVEFMEKVAKTVETEELTVEERNLLSVAYKNVIGARRASWRIISS
IEQKEDSRGNSDHVSIIKDYRGKIETELSKICDGILNLLEAHLIPAASLAESKVFYLKMKGDYHRYLAEF
KTGAERKEAAESTLVAYKSAQDIALADLAPTHPIRLGLALNFSVFYYEILNSSDRACSLAKQAFDEAISE
LDTLGEESYKDSTLIMQLLRDNLTLWTSDLNDEAGDDIKEAPKEVQKVDEQAQPPPSQ

**LAMBDA**

>gi|1549404|gb|AAB08482.1| GF14 lambda [Arabidopsis thaliana]
MAATLGRDQYVYMAKLAEQAERYEEMVQFMEQLVTGATPAEELTVEERNLLSVAYKNVIGSLRAAWRIVS
SIEQKEESRKNDEHVSLVKDYRSKVESELSSVCSGILKLLDSHLIPSAGASESKVFYLKMKGDYHRYMAE
FKSGDERKTAAEDTMLAYKAAQDIAAADMAPTHPIRLGLALNFSVFYYEILNSSDKACNMAKQAFEEAIA
ELDTLGEESYKDSTLIMQLLRDNLTLWTSDMQEQMDEA

**NU**

>gi|1531631|gb|AAB49335.1| GF14 nu [Arabidopsis thaliana]
MSSSREENVYLAKLAEQAERYEEMVEFMEKVAKTVDTDELTVEERNLLSVAYKNVIGARRASWRIISSIE
QKEESRGNDDHVSIIKDYRGKIETELSKICDGILNLLDSHLVPTASLAESKVFYLKMKGDYHRYLAEFKT
GAERKEAAESTLVAYKSAQDIALADLAPTHPIRLGLALNFSVFYYEILNSPDRACSLAKQAFDEAISELD
TLGEESYKDSTLIMQLLRDNLTLWNSDINDEAGGDEIKEASKHEPEEGKPAETGQ

**KAPPA**

>gi|17352487|gb|AAA79700.2| GF14 Kappa isoform [Arabidopsis thaliana]
MATTLSRDQYVYMAKLAEQAERYEEMVQFMEQLVSGATPAGELTVEERNLLSGAYKNVIGSLRAAWRIVS
SLEQKEESRKNEEHVSLVKDYRSKVETELSSICSGILRLLDSHLIPSATARESKVFYLKMKGDYHRYLAE
FKSGDERKTAAEDAMIAYKAAQDVAVADLAPTHPIRLGLALNFSVFYYEILNSSEKACSMAKQAFEEAIA
ELDTLGEESYKDSTLIMQLLRDNLTLWTSDMQEQMDEA

**MU**

>gi|78499631|gb|AAB49334.2| GF14 mu [Arabidopsis thaliana]
MGSGKERDTFVYLAKLSEQAERYEEMVESMKSVAKLNVDLTVEERNLLSVGYKNVIGSRRASWRIFSSIE
QKEAVKGNDVNVKRIKEYMEKVELELSNICIDIMSVLDEHLIPSASEGESTVFFNKMKGDYYRYLAEFKS
GNERKEAADQSLKAYEIATTAAEAKLPPTHPIRLGLALNFSVFYYEIMNAPERACHLAKQAFDEAISELD
TLNEESYKDSTLIMQLLRDNLTLWTSDISEEGGDDAHKTNGSAKPGAGGDDAE

**EPSILON**

>gi|1022778|gb|AAA79699.1| GF14 epsilon isoform [Arabidopsis thaliana]
MENEREKQVYLAKLSEQTERYDEMVEAMKKVAQLDVELTVEERNLVSVGYKNVIGARRASWRILSSIEQK
EESKGNDENVKRLKNYRKRVEDELAKVCNDILSVIDKHLIPSSNAVESTVFFYKMKGDYYRYLAEFSSGA
ERKEAADQSLEAYKAAVAAAENGLAPTHPVRLGLALNFSVFYYEILNSPESACQLAKQAFDDAIAELDSL
NEESYKDSTLIMQLLRDNLTLWTSDLNEEGDERTKGADEPQDEN

**OMICRON**

>gi|12044389|gb|AAG47840.1| 14-3-3 protein GF14omicron [Arabidopsis thaliana]
MENERAKQVYLAKLNEQAERYDEMVEAMKKVAALDVELTIEERNLLSVGYKNVIGARRASWRILSSIEQK
EESKGNEQNAKRIKDYRTKVEEELSKICYDILAVIDKHLVPFATSGESTVFYYKMKGDYFRYLAEFKSGA
DREEAADLSLKAYEAATSSASTELSTTHPIRLGLALNFSVFYYEILNSPERACHLAKRAFDEAIAELDSL
NEDSYKDSTLIMQLLRDNLTLWTSDLEEGGK

**IOTA**

>gi|12963453|gb|AAK11271.1|AF335544_1 14-3-3 protein GF14iota [Arabidopsis thaliana]
MSSSGSDKERETFVYMAKLSEQAERYDEMVETMKKVARVNSELTVEERNLLSVGYKNVIGARRASWRIMS
SIEQKEESKGNESNVKQIKGYRQKVEDELANICQDILTIIDQHLIPHATSGEATVFYYKMKGDYYRYLAE
FKTEQERKEAAEQSLKGYEAATQAASTELPSTHPIRLGLALNFSVFYYEIMNSPERACHLAKQAFDEAIA
ELDTLSEESYKDSTLIMQLLRDNLTLWTSDLPEDGGEDNIKTEESKQEQAKPADATEN

**PI**

>gi|332197962|gb|AEE36083.1| general regulatory factor 13 [Arabidopsis thaliana]
MENEREKLIYLAKLGCQAGRYDDVMKSMRKVCELDIELSEEERDLLTTGYKNVMEAKRVSLRVISSIEKM
EDSKGNDQNVKLIKGQQEMVKYEFFNVCNDILSLIDSHLIPSTTTNVESIVLFNRVKGDYFRYMAEFGSD
AERKENADNSLDAYKVAMEMAENSLAPTNMVRLGLALNFSIFNYEIHKSIESACKLVKKAYDEAITELDG
LDKNICEESMYIIEMLKYNLSTWTSGDGNGNKTDG

#### 3. *Homo sapiens* 14-3-3 protein gene family

#### Source: UniProt Database

Bateman, A. (2019) UniProt: A worldwide hub of protein knowledge. *Nucleic Acids Res.* **47**, D506–D515

**BETA/ALPHA**

>sp|P31946|1433B_HUMAN 14-3-3 protein beta/alpha OS=Homo sapiens GN=YWHAB PE=1 SV=3
MTMDKSELVQKAKLAEQAERYDDMAAAMKAVTEQGHELSNEERNLLSVAYKNVVGARRSS
WRVISSIEQKTERNEKKQQMGKEYREKIEAELQDICNDVLELLDKYLIPNATQPESKVFY
LKMKGDYFRYLSEVASGDNKQTTVSNSQQAYQEAFEISKKEMQPTHPIRLGLALNFSVFY
YEILNSPEKACSLAKTAFDEAIAELDTLNEESYKDSTLIMQLLRDNLTLWTSENQGDEGD
AGEGEN

**ZETA/DELTA**

>sp|P63104|1433Z_HUMAN 14-3-3 protein zeta/delta OS=Homo sapiens GN=YWHAZ PE=1 SV=1
MDKNELVQKAKLAEQAERYDDMAACMKSVTEQGAELSNEERNLLSVAYKNVVGARRSSWR
VVSSIEQKTEGAEKKQQMAREYREKIETELRDICNDVLSLLEKFLIPNASQAESKVFYLK
MKGDYYRYLAEVAAGDDKKGIVDQSQQAYQEAFEISKKEMQPTHPIRLGLALNFSVFYYE
ILNSPEKACSLAKTAFDEAIAELDTLSEESYKDSTLIMQLLRDNLTLWTSDTQGDEAEAG
EGGEN

**EPSILON**

>sp|P62258|1433E_HUMAN 14-3-3 protein epsilon OS=Homo sapiens GN=YWHAE PE=1 SV=1
MDDREDLVYQAKLAEQAERYDEMVESMKKVAGMDVELTVEERNLLSVAYKNVIGARRASW
RIISSIEQKEENKGGEDKLKMIREYRQMVETELKLICCDILDVLDKHLIPAANTGESKVF
YYKMKGDYHRYLAEFATGNDRKEAAENSLVAYKAASDIAMTELPPTHPIRLGLALNFSVF
YYEILNSPDRACRLAKAAFDDAIAELDTLSEESYKDSTLIMQLLRDNLTLWTSDMQGDGE
EQNKEALQDVEDENQ

**THETA**

>sp|P27348|1433T_HUMAN 14-3-3 protein theta OS=Homo sapiens GN=YWHAQ PE=1 SV=1
MEKTELIQKAKLAEQAERYDDMATCMKAVTEQGAELSNEERNLLSVAYKNVVGGRRSAWR
VISSIEQKTDTSDKKLQLIKDYREKVESELRSICTTVLELLDKYLIANATNPESKVFYLK
MKGDYFRYLAEVACGDDRKQTIDNSQGAYQEAFDISKKEMQPTHPIRLGLALNFSVFYYE
ILNNPELACTLAKTAFDEAIAELDTLNEDSYKDSTLIMQLLRDNLTLWTSDSAGEECDAA
EGAEN

**GAMMA**

>sp|P61981|1433G_HUMAN 14-3-3 protein gamma OS=Homo sapiens GN=YWHAG PE=1 SV=2
MVDREQLVQKARLAEQAERYDDMAAAMKNVTELNEPLSNEERNLLSVAYKNVVGARRSSW
RVISSIEQKTSADGNEKKIEMVRAYREKIEKELEAVCQDVLSLLDNYLIKNCSETQYESK
VFYLKMKGDYYRYLAEVATGEKRATVVESSEKAYSEAHEISKEHMQPTHPIRLGLALNYS
VFYYEIQNAPEQACHLAKTAFDDAIAELDTLNEDSYKDSTLIMQLLRDNLTLWTSDQQDD
DGGEGNN

**SIGMA**

>sp|P31947|1433S_HUMAN 14-3-3 protein sigma OS=Homo sapiens GN=SFN PE=1 SV=1
MERASLIQKAKLAEQAERYEDMAAFMKGAVEKGEELSCEERNLLSVAYKNVVGGQRAAWR
VLSSIEQKSNEEGSEEKGPEVREYREKVETELQGVCDTVLGLLDSHLIKEAGDAESRVFY
LKMKGDYYRYLAEVATGDDKKRIIDSARSAYQEAMDISKKEMPPTNPIRLGLALNFSVFH
YEIANSPEEAISLAKTTFDEAMADLHTLSEDSYKDSTLIMQLLRDNLTLWTADNAGEEGG
EAPQEPQS

**ETA**

>sp|Q04917|1433F_HUMAN 14-3-3 protein eta OS=Homo sapiens GN=YWHAH PE=1 SV=4
MGDREQLLQRARLAEQAERYDDMASAMKAVTELNEPLSNEDRNLLSVAYKNVVGARRSSW
RVISSIEQKTMADGNEKKLEKVKAYREKIEKELETVCNDVLSLLDKFLIKNCNDFQYESK
VFYLKMKGDYYRYLAEVASGEKKNSVVEASEAAYKEAFEISKEQMQPTHPIRLGLALNFS
VFYYEIQNAPEQACLLAKQAFDDAIAELDTLNEDSYKDSTLIMQLLRDNLTLWTSDQQDE
EAGEGN

#### 4. *Saccharomyces cerevisiae* 14-3-3 proteins

Van Heusden, G. P. H., Griffiths, D. J. F., Ford, J. C., Chin- A- Woeng, T. F. C., Schrader, P. A. T., Carr, A. M., and Steensma, H. Y. (1995) The 14- 3- 3 Proteins Encoded by the BMH1 and BMH2 Genes are Essential in the Yeast Saccharomyces cerevisiae and Can be Replaced by a Plant Homologue. *Eur. J. Biochem.* **229**, 45–53

**BMH1**

>sp|P29311|BMH1_YEAST Protein BMH1 OS=Saccharomyces cerevisiae (strain ATCC 204508 / S288c) GN=BMH1 PE=1 SV=4
MSTSREDSVYLAKLAEQAERYEEMVENMKTVASSGQELSVEERNLLSVAYKNVIGARRAS
WRIVSSIEQKEESKEKSEHQVELICSYRSKIETELTKISDDILSVLDSHLIPSATTGESK
VFYYKMKGDYHRYLAEFSSGDAREKATNASLEAYKTASEIATTELPPTHPIRLGLALNFS
VFYYEIQNSPDKACHLAKQAFDDAIAELDTLSEESYKDSTLIMQLLRDNLTLWTSDMSES
GQAEDQQQQQQHQQQQPPAAAEGEAPK

**BMH2**

>sp|P34730|BMH2_YEAST Protein BMH2 OS=Saccharomyces cerevisiae (strain ATCC 204508 / S288c) GN=BMH2 PE=1 SV=3
MSQTREDSVYLAKLAEQAERYEEMVENMKAVASSGQELSVEERNLLSVAYKNVIGARRAS
WRIVSSIEQKEESKEKSEHQVELIRSYRSKIETELTKISDDILSVLDSHLIPSATTGESK
VFYYKMKGDYHRYLAEFSSGDAREKATNSSLEAYKTASEIATTELPPTHPIRLGLALNFS
VFYYEIQNSPDKACHLAKQAFDDAIAELDTLSEESYKDSTLIMQLLRDNLTLWTSDISES
GQEDQQQQQQQQQQQQQQQQAPAEQTQGEPTK

#### 5. *Drosophila melanogaster* 14-3-3 proteins

**Source: UniProt Database**

Bateman, A. (2019) UniProt: A worldwide hub of protein knowledge. *Nucleic Acids Res.* **47**, D506–D515

**EPSILON**

>sp|P92177|1433E_DROME 14-3-3 protein epsilon OS=Drosophila melanogaster GN=14-3-3epsilon PE=1 SV=2

MTERENNVYKAKLAEQAERYDEMVEAMKKVASMDVELTVEERNLLSVAYKNVIGARRASW
RIITSIEQKEENKGAEEKLEMIKTYRGQVEKELRDICSDILNVLEKHLIPCATSGESKVF
YYKMKGDYHRYLAEFATGSDRKDAAENSLIAYKAASDIAMNDLPPTHPIRLGLALNFSVF
YYEILNSPDRACRLAKAAFDDAIAELDTLSEESYKDSTLIMQLLRDNLTLWTSDMQAEEV
DPNAGDGEPKEQIQDVEDQDVS

**ZETA**

>sp|P29310|1433Z_DROME 14-3-3 protein zeta OS=Drosophila melanogaster GN=14-3-3zeta PE=1 SV=1
MSTVDKEELVQKAKLAEQSERYDDMAQAMKSVTETGVELSNEERNLLSVAYKNVVGARRS
SWRVISSIEQKTEASARKQQLAREYRERVEKELREICYEVLGLLDKYLIPKASNPESKVF
YLKMKGDYYRYLAEVATGDARNTVVDDSQTAYQDAFDISKGKMQPTHPIRLGLALNFSVF
YYEILNSPDKACQLAKQAFDDAIAELDTLNEDSYKDSTLIMQLLRDNLTLWTSDTQGDEA
EPQEGGDN

#### 6. *Caenorhabditis elegans* 14-3-3 proteins

#### Source: UniProt Database

Bateman, A. (2019) UniProt: A worldwide hub of protein knowledge. *Nucleic Acids Res.* **47**, D506–D515

**14-3-3-like protein 1 (PAR-5)**

>sp|P41932|14331_CAEEL 14-3-3-like protein 1 OS=Caenorhabditis elegans GN=par-5 PE=1 SV=2
MSDTVEELVQRAKLAEQAERYDDMAAAMKKVTEQGQELSNEERNLLSVAYKNVVGARRSS
WRVISSIEQKTEGSEKKQQLAKEYRVKVEQELNDICQDVLKLLDEFLIVKAGAAESKVFY
LKMKGDYYRYLAEVASEDRAAVVEKSQKAYQEALDIAKDKMQPTHPIRLGLALNFSVFYY
EILNTPEHACQLAKQAFDDAIAELDTLNEDSYKDSTLIMQLLRDNLTLWTSDVGAEDQEQ
EGNQEAGN

**14-3-3-like protein 2 (FTT-2)**

>sp|Q20655|14332_CAEEL 14-3-3-like protein 2 OS=Caenorhabditis elegans GN=ftt-2 PE=1 SV=1
MSDGKEELVNRAKLAEQAERYDDMAASMKKVTELGAELSNEERNLLSVAYKNVVGARRSS
WRVISSIEQKTEGSEKKQQMAKEYREKVEKELRDICQDVLNLLDKFLIPKAGAAESKVFY
LKMKGDYYRYLAEVASGDDRNSVVEKSQQSYQEAFDIAKDKMQPTHPIRLGLALNFSVFF
YEILNAPDKACQLAKQAFDDAIAELDTLNEDSYKDSTLIMQLLRDNLTLWTSDAATDDTD
ANETEGGN

#### 7. *Solanum lycopersicum* 14-3-3 protein gene family

Xu, W. F., and Shi, W. M. (2006) Expression profiling of the 14-3-3 gene family in response to salt stress and potassium and iron deficiencies in young tomato (Solanum lycopersicum) roots: Analysis by real-time RT-PCR. *Ann. Bot.* **98**, 965–974

**TFT1**

>gi|802083923|ref|NP_001234107.2| 14-3-3 family protein [Solanum lycopersicum]
MASPREENVYMAKLAEQAERYEEMVEFMEKVVAALNGEELTVEERNLLSVAYKNVIGARRASWRIISSIE
QKEESRGNEDHVASIKKYRSQIENELTSICNGILKLLDSKLIGSAATGDSKVFYLKMKGDYYRYLAEFKT
GTERKEAAENTLSAYKSAQDIANGELAPTHPIRLGLALNFSVFYYEILNSPDRACNLAKQAFDEAIAELD
TLGEESYKDSTLIMQLLRDNLTLWTSDMQDDGTDEIKEPSKADNE

**TFT2**

>gi|823683788|ref|NP_001296299.1| 14-3-3 protein 2 [Solanum lycopersicum]
MAREENVYMAKLAEQAERYEEMVQFMEKVSTSLGSEELTVEERNLLSVAYKNVIGARRASWRIISSIEQK
EESRGNEEHVKCIKEYRSKIESELSDICDGILKLLDSNLIPSASNGDSKVFYLKMKGDYHRYLAEFKTGA

ERKEAAESTLSAYKAAQDIANTELAPTHPIRLGLALNFSVFYYEILNSPDRACNLAKQAFDEAIAELDTL
GEESYKDSTLIMQLLRDNLTLWTSDMQDDGADEIKETKNDNEQQ

**TFT3**

>gi|927442685|ref|NP_001234677.2| 14-3-3 protein 3 [Solanum lycopersicum]
MAVAPTAREENVYMAKLAEQAERYEEMVEFMEKVSNSLGSEELTVEERNLLSVAYKNVIGARRASWRIIS
SIEQKEESRGNEEHVNSIREYRSKIENELSKICDGILKLLDSKLIPSATSGDSKVFYLKMKGDYHRYLAE
FKTGAERKEAAESTLTAYKAAQDIASAELAPTHPIRLGLALNFSVFYYEILNSPDRACNLAKQAFDEAIA
ELDTLGEESYKDSTLIMQLLRDNLTLWTSDMQDDGADEIKEDPKPEEKN

**TFT4**

>gi|350536755|ref|NP_001234007.1| 14-3-3 protein 4 [Solanum lycopersicum]
MADSSREENVYLAKLAEQAERYEEMIEFMEKVAKTADVEELTVEERNLLSVAYKNVIGARRASWRIISSI
EQKEESRGNEDHVNTIKEYRSKIEAELSKICDGILSLLESNLIPSASTAESKVFYLKMKGDYHRYLAEFK
TGTERKEAAENTLLAYKSAQDIALAELAPTHPIRLGLALNFSVFYYEILNSPDRACNLAKQAFDEAISEL
DTLGEESYKDSTLIMQLLRDNLTLWTSDNADDVGDDIKEASKPESGEGQQ

**TFT5**

>gi|3023182|sp|P93210.1|14335_SOLLC RecName: Full=14-3-3 protein 5
MASPREENVYMANVADEAERYEEMVEFMEKVVAALNGEELTVEERNLLSVAYKNVIGARRASWRIISSIE
QKEESRGNEDHVASIKKYRSQIENELTSICNGILKLLDSKLIGSAATGDSKVFYLKMKGDYYRYLAEFKT
GTERKEAAENTLSAYKSAQDIANGELAPTHPIRLGLALNFSVFYYEILNSPDRACNLAKQAFDEAIAELD
TLGEESYKDSTLIMQLLRDNLTLWTSDMQDDGTDEIKEPSKADNE

**TFT6**

>gi|350538649|ref|NP_001234097.1| 14-3-3 protein 6 [Solanum lycopersicum]
MASPREENVYMAKLAEQAERYEEMVEFMEKVVAAADGAEELTVEERNLLSVAYKNVIGARRASWRIISSI
EQKEESRGNEDHVASIKEYRSKIESELTSICNGILKLLDSKLIGSAATGDSKVFYLKMKGDYHRYLAEFK
TGAERKEAAENTLSAYKAAQDIANAELAPTHPIRLGLALNFSVFYYEILNSPDRACNLAKQAFDEAIAEL
DTLGEESYKDSTLIMQLLRDNLTLWTSDMQDDGTDEIKEATPKPDDNE

**TFT7**

>gi|350539221|ref|NP_001234637.1| 14-3-3 protein 7 [Solanum lycopersicum]
MEKEREKQVYLARLAEQAERYDEMVEAMKAIAKMDVELTVEERNLVSVGYKNVIGARRASWRILSSIEQK
EESKGHEQNVKRIKTYRQRVEDELTKICSDILSVIDEHLVPSSTTGESTVFYYKMKGDYYRYLAEFKAGD
DRKEASEQSLKAYEAATATASSDLAPTHPIRLGLALNFSVFYYEILNSPERACHLAKQAFDEAIAELDSL
SEESYKDSTLIMQLLRDNLTLWTSDLEEGGEHSKGDERQGEN

**TFT8**

>gi|350539761|ref|NP_001234267.1| 14-3-3 protein 8 [Solanum lycopersicum]
MASSKERESLVYIARLAEQAERYDEMVDAMKNVANLDVELTVEERNLLSVGYKNVVGSRRASWRILSSIE
QKEDARGNEQNVKRIQGYRQKVESELTDICNNIMTVIDEHLIPSCTAGESTVFYYKMKGDYYRYLAEFKT
GDDKKEVSDLSLKAYQTATTTAEAELPITHPIRLGLALNFSVFYYEIMNSPERACQLAKQVFDEAISELD
SLNEDNYKDGTLILQLLRDNLTLWTSDIPEDGEEAPKGDAANKVGAGEDAE

**TFT9**

>gi|350536935|ref|NP_001234272.1| 14-3-3 protein 9 [Solanum lycopersicum]
MASSKERENFVYVAKLAEQAERYDEMVEAMKNVANMDVELTVEERNLLSVGYKNVVGSRRASWRILSSIE
QKEESRGNEQNVKRIKEYLQKVESELTNICNDIMVVIDQHLIPSCSAGESTVFYHKMKGDYYRYLAEFKA
GNDKKEVAELSLKAYQAATTAAEAELAPTHPIRLGLALNFSVFYYEIMNSPERACHLAKQAFDEAISELD
SLNEDSYKDSTLIMQLLRDNLTLWTSDLPEDAEDAQKGDATNKAGGGEDAE

**TFT10**

>gi|350539807|ref|NP_001234278.1| 14-3-3 protein 10 [Solanum lycopersicum]
MAALIPENLSREQCLYLAKLAEQAERYEEMVQFMDKLVLNSTPAGELTVEERNLLSVAYKNVIGSLRAAW
RIVSSIEQKEESRKNEEHVHLVKEYRGKVENELSQVCAGILKLLESNLVPSATTSESKVFYLKMKGDYYR
YLAEFKIGDERKQAAEDTMNSYKAAQEIALTDLPPTHPIRLGLALNFSVFYFEILNSSDKACSMAKQAFE
EAIAELDTLGEESYKDSTLIMQLLRDNLTLWTSDAQDQLDES

#### 8. Other 14-3-3 Family Members

Tian, F., Wang, T., Xie, Y., Zhang, J., and Hu, J. (2015) Genome-wide identification, classification, and expression analysis of 14-3-3 gene family in Populus. *PLoS One*. **10**, e0123225.

##### *Populus trichocarpa*

```
>gi|566146429|ref|XP_006368230.1| 14-3-3 brain family protein [Populus
trichocarpa]
MDVRQNSVYSAKLAEQAKRYNEMLDHMKYIAKLDVELTTEERNLLRIGCKNVMGPRRESWRMLSSIEEKE
QAKGNQVNAKRIEEYRRKIESELTSICNDIIQLIDDHLLPSTSQCESCVFYHRMKGDYYRYLAEFKVGIE
MEKAASESMKAYDIGIKAASKLAPTNLVRLSLALNFSVLLYDIMKFPEKAFFHAKNAYDEAIPILDNLNK
ESQKDSMLILEILLDNVRLWSYDILEN
>gi|224063273|ref|XP_002301071.1| 14-3-3 protein 32kDa endonuclease [Populus
trichocarpa]
MAVTPSAREENVYMAKLAEQAERYEEMVEYMEKVSASLENEELTVEERNLLSVAYKNVIGARRASWRIIS
SIEQKEESRGNEDHVSVIRDYRAKIETELSSICDGILKLLDSRLIPTASAGDSKVFYLKMKGDYHRYLAE
FKTGAERKEAAESTLTAYKAAQDIANAELAPTHPIRLGLALNFSVFYYEILNSPDRACSLAKQAFDEAIA
ELDTLGEESYKDSTLIMQLLRDNLTLWTSDMQDDGADEIKEAAPKPGDEQQ
>gi|566213236|ref|XP_006373446.1| 14-3-3 family protein [Populus trichocarpa]
MSPTEPSREESVYMAKLAEQAERYEEMVEFMEKVAKTVDNEELTMEERNLLSVAYKNVIGARRASWRIIS
SIEQKEESRGNEDHVTIIKEYRGKIEAELSKICDGILSLLETHLVPSASAAESKVFYLKMKGDYHRYLAE
FKTGAERKEAAESTLLSYKSAQDIALSELAPTHPIRLGLALNFSVFYYEILNSPDRACSLAKQAFDEAIS
ELDTLGEESYKDSTLIMQLLRDNLTLWTSDITDEAGDEIKDASKRESGDGPQ
>gi|224084622|ref|XP_002307363.1| 14-3-3 protein 32kDa endonuclease [Populus
trichocarpa]
MAATPSAREENVYMAKLAEQAERYEEMVEYMEKVSASIDNEELTVEERNLLSVAYKNVIGARRASWRIIS
SIEQKEESRGNEDHVSVIRDYRAKIETELSSICDGILKLLDTRLIPTASSGDSKVFYLKMKGDYHRYLAE
FKTGAERKEAAESTLTAYKSAQDIANAELAPTHPIRLGLALNFSVFYYEILNSPDRACNLAKQAFDEAIA
ELDTLGEESYKDSTLIMQLLRDNLTLWTSDMQDDAADEIKEAAPKTGDEQ
>gi|224114812|ref|XP_002316863.1| 14-3-3 brain family protein [Populus
trichocarpa]
MDSSKDRENFVYVAKLAEQAERYDEMVDAMKKVAKLDVELTVEERNLLSVGYKNVIGARRASWRILSSIE
QKEESKGNETNVKRIKEYRKKVEAELTGVCNDIMTVIDEHLIPSSIPGESSVFYHKMKGDYYRYLAEFKS
GNERKEAADQSLKAYETATSTAARDLSPTHPIRLGLALNFSVFYYEIMNSPERACHLAKQSFDEAISELD
TLSEESYKDSTLIMQLLRDNLTLWTSDIPEDGEDQKMETSARAGGGEDAE
```

##### *Oryza sativa*

```
>gi|115435206|ref|NP_001042361.1| Os01g0209200 [Oryza sativa Japonica Group]
MAPSDDLVYMAKLAEQAERYDEMVEAMNSVAKLDEGLTKEERNLLSVGYKNLIGAKRAAMRIIGSIELKE
ETKGKESHVRQTAEYRRKVEAEMDKICCDVINIIDKYLIPHSSGAESSVFYYKMKGDYYRYLAEFKTGTE
KIEVSELSLNAYETASKTAQTDLTPTDPIRLGLALNISVFYCEIMNSPDKACQLAKNAFDEAVAELPSLS
EENYKDSTLIMQLLRDNLALWNSDMADDADDIRERTDTTGAKGDPAA
>gi|115446909|ref|NP_001047234.1| Os02g0580300 [Oryza sativa Japonica Group]
MSQPAELSREENVYMAKLAEQAERYEEMVEFMEKVAKTVDSEELTVEERNLLSVAYKNVIGARRASWRII
SSIEQKEESRGNEDRCTLIKEYRGKIETELSKICDGILKLLDSHLVPSSTAPESKVFYLKMKGDYYRYLA
EFKTGAERKDAAENTMVAYKAAQDIALAELPPTHPIRLGLALNFSVFYYEILNSPDRACNLAKQAFDEAI
SELDTLSEESYKDSTLIMQLLRDNLTLWTSDISEDAAEEIKEAPKGESGDGQ
>gi|115454901|ref|NP_001051051.1| Os03g0710800 [Oryza sativa Japonica Group]
MSPAEASREENVYMAKLAEQAERYEEMVEFMEKVAKTTDVGELTVEERNLLSVAYKNVIGARRASWRIIS
SIEQKEESRGNEAYVASIKEYRSRIETELSKICDGILKLLDSHLVPSATAAESKVFYLKMKGDYHRYLAE
```

FKSGAERKEAAENTLVAYKSAQDIALADLPTTHPIRLGLALNFSVFYYEILNSPDRACNLAKQAFDDAIA
ELDTLGEESYKDSTLIMQLLRDNLTLWTSDNAEDGGDEIKEAAKPEGEGH
>gi|115458806|ref|NP_001053003.1| Os04g0462500 [Oryza sativa Japonica Group]
MSAQAELSREENVYMAKLAEQAERYEEMVEFMEKVAKTVDSEELTVEERNLLSVAYKNVIGARRASWRII
SSIEQKEESRGNEDRVTLIKDYRGKIETELTKICDGILKLLESHLVPSSTAPESKVFYLKMKGDYYRYLA
EFKTGAERKDAAENTMVAYKAAQDIALAELPPTHPIRLGLALNFSVFYYEILNSPDRACNLAKQAFDEAI
SELDTLSEESYKDSTLIMQLLRDNLTLWTSDISEDTAEEIREAPKRDSSEGQ
>gi|115476520|ref|NP_001061856.1| Os08g0430500 [Oryza sativa Japonica Group]
MSREENVYMAKLAEQAERYEEMVEYMEKVAKTVDVEELTVEERNLLSVAYKNVIGARRASWRIVSSIEQK
EEGRGNEEHVTLIKEYRGKIEAELSKICDGILKLLDSHLVPSSTAAESKVFYLKMKGDYHRYLAEFKTGA
ERKEAAESTMVAYKAAQDIALADLAPTHPIRLGLALNFSVFYYEILNSPDKACNLAKQAFDEAISELDTL
GEESYKDSTLIMQLLRDNLTLWTSDLTEDGGDEVKEASKGDACEGQ
>gi|115476928|ref|NP_001062060.1| Os08g0480800 [Oryza sativa Japonica Group]
MAAAAGGGTREEMVYMAKLAEQAERYEEMVEFMEKVVTAAAAGGGGELTVEERNLLSVAYKNVIGARRAS
WRIVSSIEQKEEGRGAAGHAAAARSYRARVEAELSNICAGILRLLDERLVPAAAAVDAKVFYLKMKGDYH
RYLAEFKTGAERKDAADATLAAYQAAQDIAMKELSPTHPIRLGLALNFSVFYYEILNSPDRACTLAKQAF
DEAISELDTLGEESYKDSTLIMQLLRDNLTLWTSDMQDDGGDEMRDATKPEDEH
>gi|297611974|ref|NP_001068067.2| Os11g0546900 [Oryza sativa Japonica Group]
MSPAEPTREESVYKAKLAEQAERYEEMVEYMERVARAAGGASGGEELTVEERNLLSVAYKNVIGARRASW
RIISSIEQKEEGRGNDAHAATIRSYRGKIEAELARICDGILALLDSHLVPSAGAAESKVFYLKMKGDYHR
YLAEFKSGDERKQAAESTMNAYKAAQDIALADLAPTHPIRLGLALNFSVFYYEILNSPDRACNLAKQAFD
EAISELDSLGEESYKDSTLIMQLLRDNLTLWTSDANDDGGDEIKEAAAPKEPGDQ

#### *Glycine max*

>gi|351726936|ref|NP_001238423.1| 14-3-3 protein SGF14e [Glycine max]
MSAEKERETQVYLAKLAEQAERYEEMVECMKKVAKLDLDLTVEERNLLSVGYKNVIGARRASWRIMSSIE
QKEESKGNEHNVKLIKSYCQKVEEELSKICGDILTIIDQHLIPSSASAEASVFYYKMKGDYFRYLAEFKT
DQERKEAAEQSLKGYEAASATANTDLPSTHPIRLGLALNFSVFYYEIMNSPERACHLAKQAFDEAIAELD
TLSEESYKDSTLIMQLLRDNLTLWTSDLPEDGGEDSIKAEETKPSEPEH
>gi|351726463|ref|NP_001238407.1| 14-3-3 protein SGF14f [Glycine max]
MSVEKERETQVYLAKLAEQAERYEEMVECMKKVAKLDLDLTVEERNLLSVGYKNVIGARRASWRIMSSIE
QKEESKGNEHNVKLIKSYCQKVEEELSKICGDILTIIDQHLIPSSGSAEASVFYYKMKGDYFRYLAEFKT
DQERKEAAEQSLKGYEAASATANTDLPSTHPIRLGLALNFSVFYYEIMNSPERACHLAKQAFDEAIAELD
TLSEESYKDSTLIMQLLRDNLTLWTSDLPEDGGEDNIKAEEEAKPSEPEH
>gi|351724401|ref|NP_001238592.1| 14-3-3 protein SGF14n [Glycine max]
MTQPAMATFSKERENFVYVAKLAEQAERYDEMVDAMKKVAKLDVELSVEERNLFSVGYKNVVGSRRASWR
ILSSIEQKEESKGNELHVKRIRDYRNKVELELSNICSDIMIVLDEHLIPSTNIAESTVFYYKMKGDYYRY
LAEFKAGNEKKEVADQSLKAYETASTTAESELQPTHPIRLGLALNFSVFYYEIMNSPERACHLAKQAFDD
AVSDLDTLNEDSYKDSTLIMQLLRDNLTLWTSDIPEEGEDQKMESTTRGEDELGR

#### *Medicago trucatula*

>gi|357452267|ref|XP_003596410.1| general regulatory factor 2 [Medicago truncatula]
MASSTNVRENFVYVAKLAEQAERYDEMVEAMKKLAKMDVELSVEERNLFSVGYKNVVGSRRASWRILSSI
EQKEESKGNELNVKRIKEYRQKVEVELSSICNDIMIIIDEHLIPSTNIAESTVFYYKMKGDYYRYLAEFK
AGDEKKEVADLSLKAYQTASATAENELQPTHPIRLGLALNFSVFYYEIMNSPERACHLAKQAFDDGVSEL
DTLNEDSYKDSTLIMQLLRDNLTLWTSDIPEDGEDQKMESATKSGQDEDELGR
>gi|922376198|ref|XP_003602823.2| general regulatory factor 2 [Medicago truncatula]
MAAAHSPREENVYMAKLAEQAERYEEMVEFMEKVSANADNEELTVEERNLLSVAYKNVIGARRASWRIIS

SIEQKEESRGNEDHVSVIRDYRSKIESELSNICDGILKLLDSRLIPSAASGDSKVFYLKMKGDYHRYLAE
FKTGAERKEAAESTLAAYKSAQDIANAELPPTHPIRLGLALNFSVFYYEILNSPDRACNLAKQAFDEAIA
ELDTLGEESYKDSTLIMQLLRDNLTLWTSDMQDDGADEIKEAAPKPDEQQ
>gi|357465319|ref|XP_003602941.1| general regulatory factor 2 [Medicago truncatula]
MGGAIPENLNREQYVYLAKLAEQAERYEEMVSFMQKLVVGSTPSSELSVEERNLISVAYKNVIGSLRAAW
RIVSSIEQKEEGRKNEDHVVLVKDYRSKVESELTNVCGSILELLDSNLIPSASSSESKVFYYKMKGDYHR
YLAEFKIGDEKKSAAEDTMLSYKAAQDIAAADLPSTHPIRLGLALNFSVFYYEILNQSDKACDMAKQAFE
EAIAELDTLGEESYKDSTLIMQLLRDNLTLWTSDVQDQLDEP
>gi|922363262|ref|XP_003607800.2| general regulatory factor 2 [Medicago truncatula]
MSTEKERETQVYMAKLAEQAERYEEMVECMKTIAKLDVELTVEERNLLSVGYKNVIGARRASWRIMSSIE
QKEEAKGNENNVKLIKSYCQKVEEELSKICSDILEIIDKHLIPSSTTGEATVFYYKMKGDYYRYLAEFKN
DQDRKEAADQSLKAYEAASATASTDLPSTHPIRLGLALNFSVFYYEIMNSPERACHLAKQAFDEAIADLD
TLSEESYKDSTLIMQLLRDNLTLWTSDLPEDGGEELKSEEVKPAEPEGEPARLTRWKQLCYDDNRRKRRR
STIYS
>gi|357489745|ref|XP_003615160.1| general regulatory factor 2 [Medicago truncatula]
MATAPTPREEFVYMAKLAEQAERYEEMVDFMEKVTAAVESEELTVEERNLLSVAYKNVIGARRASWRIIS
SIEQKEESRGNDEHVTVIRDYRSKIEAELSNICNGILKLLDSRLIPSAASGDSKVFYLKMKGDYHRYLAE
FKSGAERKDAAESTLTAYKSAQDIANSELPPTHPIRLGLALNFSVFYYEILNSPDRACGLAKQAFDEAIA
ELDTLGEESYKDSTLIMQLLRDNLTLWTSDMQDDGADEIKEAAPKGADEQ
>gi|922329887|ref|XP_003629763.2| general regulatory factor 2 [Medicago truncatula]
MASTKERENFVYIAKLAEQAERYEEMVEAMKNVAKLDVELTVEERNLLSVGYKNVVGAHRASWRILSSIE
HKEESKGYDVNVKRIKEYRHKVESELSNICSDIMSIIDDHLIPSSSAGESSVFFYKMKGDYYRYLAEFKN
GDERKEAADHSMEAYQTASTAAEGELPPTHPIRLGLALNFSVFYYEILNSPERACHLAKQAFDEAISELD
TLNEESYKDSTLIMQLLRDNLTLWTSDIPEDGDEEHKVESSGAVGGENA

### BLASTp RESULTS

1. **Homology of *Plasmodium falciparum 3D7* 14-3-3I protein with 14-3-3 proteins from plants**

**Query Sequence:** 14-3-3I protein, putative, *Plasmodium falciparum 3D7*; MAL8P1.69
**Subject:** Plants (taxid:3193)

##### Protein sequences showing homology with 14-3-3I, *Plasmodium falciparum 3D7*

```
>gi|674916968|emb|CDY16161.1| BnaA09g28950D [Brassica napus]
MSSGSDKERETFVYLAKLSEQAERYDEMVETMKKVAKVDSELTVEERNLLSVGYKNVIGARRASWRIMSSIEQKEES
KGNESNVKTIKGYRQKVEDELANICQDILSIIDQHLIPHATSGEATVFYYKMKGDYYRYLAEFKTEQERKEASEQSL
KGYEAATQAASTELPSTHPIRLGLALNFSVFYYEIMNSPERACHLAKQAFDEAIAELDTLSEESYKDSTLIMQLLRD
NLTLWTSDLPEDGGEDTKTDEPKKEEAKPAEATEVIDL
>gi|674948527|emb|CDX85023.1| BnaC05g20320D [Brassica napus]
MSSGTDKEREIFVYLAKLSEQAERYDEMVETMKKVAKVDSELTVEERNLLSVGYKNVIGARRASWRIMSSIEQKEES
KGNESNVKTIKGYRQKVEDELANICQDILSIIDQHLIPHATSGEATVFYYKMKGDYYRYLAEFKTEQERKEASEQSL
KGYEAATQAASTELPSTHPIRLGLALNFSVFYYEIMNSPERACHLAKQAFDEAIAELDTLSEESYKDSTLIMQLLRD
NLTLWTSDLPEDGGEDNAKTDEPKKEEAKPAEATEVINL
>gi|787519435|gb|AKA21457.1| 14-3-3f protein [Morus alba var. atropurpurea]
MASTKERENFVYVAKLAEQAERYDEMVDAMKKVAKLDVELTVEERNLLSVGYKNVIGARRASWRILSSIEQKEEAKG
NDQNVKRIKEYRNKVESELSDICSDIMSVVDEHLIPSCTAGESTVFFYKMKGDYYRYLAEFKAGNGKKEAADQSMKA
YETASTTAEAELPPTHPIRLGLALNFSVFYYEILNSPERACHLAKQAFDEAISELDTLSEESYKDSTLIMQLLRDNL
TLWTSDIPEDGGKNTNGSHHSILFYSILSMLFQK
>gi|672241334|gb|AIJ04693.1| 14-3-3d [Morus alba var. atropurpurea]
MSSAEKERETQVYLAKLAEQAERYEEMVESMKKLAKLDVELTVEERNLLSVGYKNVIGARRASWRIMSSIEQKEESK
GNENNVKLIKGYRQKVEDELSKICSDILTIIDKHLIPSSASGEATVFYYKMKGDYYRYLAEFKTEQDRKEAAEQSLK
GYEAASSTANAELPSTHPIRLGLALNFSVFYYEIMNSPERACHLAKQAFDEAIADLDTLSEESYKDSTLIMQLLRDN
LTLWTSDLPEDGGEDNFKGEELKPADPEH
>gi|390190105|dbj|BAM20996.1| 14-3-3 protein [Marchantia polymorpha]
MAARDELREENVYMAKLAEQAERYDEMVEAMEKVAKTVDVEELTVEERNLLSVAYKNVIGARRASWRIISSIEQKEE
SKGNDDHVAMIKDYRAKVESELSTICESILNLLDTHLIPTSTTGESKVFYLKMKGDYHRYLAEFKTGAERKEAAEST
LLAYKSAQDIALTELAPTHPIRLGLALNFSVFYYEILNSPDRACTLAKQAFDEAIAELDTLGEESYKDSTLIMQLLR
DNLTLWTSDMQEDAGDDTKEAKTEDAEDS
>gi|126508572|gb|ABO15468.1| 14-3-3 protein Lil 1433-3 [Lilium longiflorum]
MTTEKERENHVYLAKLAEQAERYDEMVESMKNVARLDLDLTVEERNLLSVGYKNVIGARRASWRIMSSIEQKEESKG
NDQNVKLIKGYRHKVEEELSKICNDILTVIDKHLIPSSGSGEATVFYYKMKGDYYRYLAEFKNEHERKEAADQSMKA
YQAASNTANTDLPSTHPIRLGLALNFSVFYYEIMNSPERACHLAKQAFDEAIAELDTLSEESYKDSTLIMQLLRDNL
TLWTSDLPEDGGDDGFKGEEAKAAGDDEH
>gi|543176570|gb|AGV54308.1| 14-3-3 protein [Phaseolus vulgaris]
MASTKERDNFVYVAKLAEQAERYEEMVDAMKNVAKLNVELTVEERNLLSVGYKNVVGARRASWRILSSIEQKEEAKG
NDVSVKRIKEYRQKVESELSNICSDIMTVIDEHLIPSSTTGEPSVFFYKMKGDYYRYLAEFKSGDERKEAADHSMKA
YQTASTTAEAELPPTHPIRLGLALNFSVFYYEILNSPERACHLAKQAFDEAIFELDTLSEESYKDSTLIMQLLRDNL
TLWTSDIPEDGEEQKVDSARATGGEEA
>gi|922376198|ref|XP_003602823.2| general regulatory factor 2 [Medicago
truncatula]
MAAAHSPREENVYMAKLAEQAERYEEMVEFMEKVSANADNEELTVEERNLLSVAYKNVIGARRASWRIISSIEQKEE
SRGNEDHVSVIRDYRSKIESELSNICDGILKLLDSRLIPSAASGDSKVFYLKMKGDYHRYLAEFKTGAERKEAAEST
LAAYKSAQDIANAELPPTHPIRLGLALNFSVFYYEILNSPDRACNLAKQAFDEAIAELDTLGEESYKDSTLIMQLLR
DNLTLWTSDMQDDGADEIKEAAPKPDEQQ
>gi|590699319|ref|XP_007045894.1| General regulatory factor 12, IOTA isoform
2 [Theobroma cacao]
```

MSNTEKERETHVYMAKLAEQAERYEEMVETMKQVAKLGCELTVEERNLLSVGYKNVIGARRASWRIMSSIEQKEESK
GNEENVKLIKGYRQKVEEELSKICTDILGIIDKHLIPSSNSGEATVFYYKMKGDYYRYVAEFKTDQERKEAAEQSLK
GYEAASAAANTDLPSTHPIRLGLALNFSVFHYEIMNSPERACHLAKQAFDEAISELDTLSEESYKDSTLIMQLLRDN
LTLWTSDLPEDGGEDSVKGEDGKSTAEAEK

#### 2. Homology of *Plasmodium falciparum 3D7* 14-3-3I protein with 14-3-3 proteins from Fungi

**Query Sequence:** 14-3-3I protein, putative, *Plasmodium falciparum 3D7*; MAL8P1.69
**Subject:** Fungi (taxid:4751)

##### Protein sequences showing homology with 14-3-3I, *Plasmodium falciparum 3D7*

>gi|907089386|gb|KNC95880.1| 14-3-3 family protein epsilon [Spizellomyces punctatus DAOM BR117]
MAEKGGREDQVYMAKLAEQAERYDEMVSYMKEVAKLGVELTVEERNLLSVAYKNVIGARRASWRIVSSIEQKEESKG
NEGQVKKIKDYRLKIETELSEVCSDILNVLDEHLIPAAEAGESKVFYYKMKGDYHRYLAEFATGEKRKEAASHAHDA
YKAATDIAQTELAPTHPIRLGLALNFSVFYYEILNSPDRACHLAKQAFDDAIAELDTLSEESYKDSTLIMQLLRDNL
TLWTSDLQDAEGDKTEEAKADEPEEAK
>gi|927373527|ref|XP_013932614.1| protein BMH2 [Ogataea parapolymorpha DL-1]
MPASREDSVYLAKLAEQAERYEEMVENMKAVASSGQELSVEERNLLSVAYKNVIGARRASWRIVSSIEQKEEAKGNE
TQVSLIREYRAKIEEELSNICEDILTVLTQHLIPTAQSGESKVFYYKMKGDYHRYLAEFAVTEKRKEAADLSLEAYK
AASEVAVTELPPTHPIRLGLALNFSVFYYEILNSPDRACHLAKQAFDDAIAELDTLSEESYKDSTLIMQLLRDNLTL
WTSDMSEAGQDEPAPEKAAEKPDDE
>gi|909128019|gb|KNE55177.1| 14-3-3 family protein epsilon [Allomyces macrogynus ATCC 38327]
MATERDNQVYMAKLSEQAERYDEMVTFMKEVAKLGVELTVEERNLLSVAYKNVIGARRASWRIVSSIEQKEENKGNV
AQVEKIKGYRQKIEAELSDVCSDILAVLDAHLIPAAEAGESKVFYYKMKGDYHRYLAEFAAGEGRKEAAAAAHDAYK
LATDIAQTELAPTHPIRLGLALNFSVFYYEILNSPDRACHLAKQAFDDAIAELDTLSEESYKDSTLIMQLLRDNLTL
WTSDLQEGDKADEAKEGEGEQA
>gi|663447431|emb|CDR45889.1| CYFA0S20e01332g1_1 [Cyberlindnera fabianii]
MSLSREDSVYLAKLAEQAERYEEMVENMKAVAGSGQELSVEERNLLSVAYKNVIGARRASWRIVSSIEQKEEAKGNE
TQVNLIREYRSKIESELTKICDDILSVLTDHLIPSAASGESKVFYYKMKGDYHRYLAEFATADRRKEAADLSLEAYK
SASDVAVTELPPTHPIRLGLALNFSVFYYEILNSPDRACHLAKQAFDDAIAELDTLSEESYKDSTLIMQLLRDNLTL
WTSDMSETGQEESAQAEEKADTKDEE
>gi|84618083|emb|CAJ16742.1| 14-3-3 protein [Rhizophagus intraradices]
MERENQTSNKKVYMAKLAEQAERYDEMVSYMKDVAKLGVELTVEERNLLSVAYKNVIGARRASWRIISSIEQKEENK
GNETQVQKIKAYRQKVETELSEVCSDILAVLDEHLIPSAEAGESKVFYYKMKGDYHRYLAEFASGEGRKEAATQAHE
AYKHATDIAQTDLAPTHPIRLGLALNFSVFYYEILNSPDRACHLAKQAFDDAIAELDTLSEESYKDSTLIMQLLRDN
LTLWTSDLQDAEGEKPEEAKQEVEAEESK
>gi|758348546|dbj|GAN09196.1| 14-3-3 family protein [Mucor ambiguus]
MSTERENNVYMAKLAEQAERYDEMVSFTKDVAKMGVELTVEERNLLSVAYKNVIGARRASWRIVSSIEQKEESKGNT
AQVEKIKTYRQKIENELQDVCKDILNVLSENLIPNAQGGESKVFYYKMMGDYHRYLAEFLTSESRKESATQAHEAYK
TATEIAQTELAPTHPIRLGLALNFSVFYYEILNSPDRACHLAKQAFDDAIAELDTLSEESYKDSTLIMQLLRDNLTL
WTSDLQEETEKSDEAKAEPVDESK
>gi|511001406|gb|EPB82885.1| 14-3-3 family protein [Mucor circinelloides f. circinelloides 1006PhL]
MAKLAEQAERYDEMVQFTKEVAKMGIELTVEERNLLSVAYKNVIGARRASWRIVSSIEQKEESKGNASQVEKIKAYR
QKIENELQGVCQDILNVLTDDLIPNAQGGESKVFYYKMKGDYHRYLAEFLTGDARKSAATEAHDAYKNATDVAQTEL
APTHPIRLGLALNFSVFYYEILNSPDRACHLAKQAFDDAIAELDTLSEESYKDSTLIMQLLRDNLTLWTSDLQDESD
KPENKPEDAEESK
>gi|661189777|emb|CDH48636.1| 14-3-3 protein [Lichtheimia corymbifera JMRC:FSU:9682]
MSNEREDKVYMAKIAEQAERYDEMVNYMKDVAKMGVDLSVEERNLLSVAYKNVIGARRASWRIVSSIEQKEESKGHE
AQVAKIKEYRNKVEKELYEVCADILALLSDYLIPSASEGEAKVFYYKMKGDYHRYVAEYATGEDRKSAASAAHEAYK

TATDVAQVDLATTHPIRLGLALNFSVFYYEILNSPDRACHLAKQAFDDAIAELDTLSEESYKDSTLIMQLLRDNLTL
WTSDLQEEGDKTDKPKEEETQE
>gi|761954227|gb|KIY73860.1| 14-3-3 protein [Cylindrobasidium torrendii FP15055 ss-10]
MPETREDSVYLAKLAEQAERYEEMVENMKRVASSDQELTVEERNLLSVAYKNVIGARRASWRIVSSIEQKEESKGNE
AQVTMIKGYREKIESELAKICEDILDVLEKHLIPSAASGESKVFYHKMMGDYHRYLAEFATGDKRKESADKSLEAYK
AASDVAVTELPPTHPIRLGLALNFSVFYYEILNSPDRACHLAKQAFDDAIAELDTLSEESYKDSTLIMQLLRDNLTL
WTSDMQDSDQKEETAEAPADE
>gi|636607341|ref|XP_008034805.1| 14-3-3 [Trametes versicolor FP-101664 SS1]
MTQTREDSVYLAKLAEQAERYEEMVENMKRVASSDQELTVEERNLLSVAYKNVIGARRASWRIVSSIEQKEESKGNE
AQVKMIKGYREKIESELAKICEDILDVLDKHLIPSAASGESKVFYHKMMGDYHRYLAEFATGDKRKESADKSLEAYK
AASDVAVTELPPTHPIRLGLALNFSVFYYEILNSPDRACHLAKQAFDDAIAELDTLSEESYKDSTLIMQLLRDNLTL
WTSDMQESDKPADKEEGAEAPAEDA
>gi|6692796|dbj|BAA89421.1| 14-3-3 [Lentinula edodes]
MPETREDSVYLAKLAEQAERYEEMVENMKRVASSDQELTVEERNLLSVAYKNVIGARRASWRIVSSIEQKEESKGNE
AQVSMIKGYREKIETELAEICEDILDVLDKHLIPSAASGESKVFYHKMMGDYHRYLAEFATGDKRKESADKSLEAYK
AASDVAVTELPPTHPIRLGLALNFSVFYYEILNSPDRACHLAKQAFDDAIAELDTLSEESYKDSTLIMQLLRDNLTL
WTSDMQDSADKPAEKDEAADAPADE
>gi|630345911|ref|XP_007862256.1| DNA damage checkpoint protein rad24 [Gloeophyllum trabeum ATCC 11539]
MSQSREDSVYLAKLAEQAERYEEMVENMKRVASSDQELTVEERNLLSVAYKNVIGARRASWRIVSSIEQKEESKGNE
AQVQMIKGYREKIESELAKICEDILDVLDKHLIPSAASGESKVFYHKMKGDYHRYLAEFATGDRRKESADKSLEAYK
AASDVAVTELPPTHPIRLGLALNFSVFYYEILNSPDRACHLAKQAFDDAIAELDTLSEESYKDSTLIMQLLRDNLTL
WTSDMQDSDKPADKEEAAEAPAEEAA
>gi|557996568|gb|EST06061.1| DNA damage checkpoint protein rad24 [Pseudozyma brasiliensis GHG001]
MPESREDSVYLAKLAEQAERYEEMVENMKRVASSDQELTVEERNLLSVAYKNVIGARRASWRIVSSIEQKEESKGNE
TQVSMIKTYREKIEAELAQICEDILDVLDKHLIPSAASGESKVFYHKMKGDYHRYLAEFATGDKRKDSADKSLEAYK
AASDVAVTELPPTHPIRLGLALNFSVFYYEILNSPDRACHLAKQAFDDAIAELDTLSEESYKDSTLIMQLLRDNLTL
WTSDMQDSEKPTEAAAEAPAAEAKGEEAA
>gi|598051181|ref|XP_007354963.1| 14-3-3 1 protein [Auricularia subglabra TFB-10046 SS5]
MSETREDSVYLAKLAEQAERYEEMVENMKRVASSDQELTVEERNLLSVAYKNVIGARRASWRIVSSIEQKEESKGNE
AQVSMIKGYREKIESELAKICEDILDVLDKHLIPSAASGESKVFYHKMMGDYHRYLAEFATGDKRKESADKSLEAYK
AASDVAVTELPPTHPIRLGLALNFSVFYYEILNSPDRACHLAKQAFDDAIAELDTLSEESYKDSTLIMQLLRDNLTL
WTSDMQDTNEKAEDKPEADVNADDA
>gi|471888554|emb|CCO32840.1| 14-3-3 protein homolog AltName: Full=Th1433 [Rhizoctonia solani AG-1 IB]
MSSSREDSVYLAKLAEQAERYEEMVENMKRVASSDQELTVEERNLLSVAYKNVIGARRASWRIVSSIEQKEESKGND
AQVTMIKGYREKIEAELAKICEDILDVLDKHLIPSAASGESKVFYHKMMGDYHRYLAEFATGDKRKESADKSLEAYK
SASDVAITELPPTHPIRLGLALNFSVFYYEILNSPDRACHLAKQAFDDAIAELDTLSEESYKDSTLIMQLLRDNLTL
WTSDMQDSADKSGDKDEAPAEADDASKA
>gi|628851406|ref|XP_007773520.1| 14-3-3 protein [Coniophora puteana RWD-64-598 SS2]
MPESREDSVYLAKLAEQAERYEEMVENMKRVASSDQELTVEERNLLSVAYKNVIGARRASWRIVSSIEQKEESKGNE
AQVAMIKSYREKIESELAKICEDILDVLDKHLIPSAASGESKVFYHKMMGDYHRYLAEFATGDKRKESADKSLEAYK
NASDVAMTELPPTHPIRLGLALNFSVFYYEILNSPDRACHLAKQAFDDAIAELDTLSEESYKDSTLIMQLLRDNLTL
WTSDMQETDTKDEPAEAPADE
>gi|906320107|emb|CEP20716.1| BMH1 [Cyberlindnera jadinii]
MSLSREDSVYLAKLAEQAERYEEMVENMKAVASSGQELSVEERNLLSVAYKNVIGARRASWRIVSSIEQKEEAKGNE
AQVALIREYRAKIETELTKICDDILSVLTNHLIPSAATGESKVFYYKMKGDYHRYLAEFATNDRRKEAADLSLEAYK
AASDVAVTELPPTHPIRLGLALNFSVFYYEILNSPDRACHLAKQAFDDAIAELDTLSEESYKDSTLIMQLLRDNLTL
WTSDMSEAGQEDAPAEEKPADAKADEE
>gi|511007036|gb|EPB88335.1| 14-3-3 family protein epsilon [Mucor circinelloides f. circinelloides 1006PhL]
MSTERENNVYMAKLAEQAERYDEMVSFTKDVAKMGVELTVEERNLLSVAYKNVIGARRASWRIVSSIEQKEESKGNT
AQVEKIKAYRQKIENELQDVCKDILQVLSENLIPNAQGGESKVFYYKMMGDYHRYLAEFLTSESRKESATQAHEAYK

TATEIAQTELAPTHPIRLGLALNFSVFYYEILNSPDRACHLAKQAFDDAIAELDTLSEESYKDSTLIMQLLRDNLTL
WTSDLQEETEKSDEAKAEPVDESK
>gi|639556755|gb|KDN34974.1| 14-3-3 protein [Rhizoctonia solani AG-8
WAC10335]
MTDTREDSVYLAKLAEQAERYEEMVENMKRVASSDQELTVEERNLLSVAYKNVIGARRASWRIVSSIEQKEESKGND
AQVTMIKGYREKIEAELAKICEDILDVLDKHLIPSAASGESKVFYHKMMGDYHRYLAEFATGDKRKQSADKSLEAYK
AASDVAVTELPPTHPIRLGLALNFSVFYYEILNSPDRACHLAKQAFDDAIAELDTLSEESYKDSTLIMQLLRDNLTL
WTSDMQDSADKSGDKDEAPAEADDAPKA
>gi|914968799|ref|XP_013240790.1| 14-3-3 protein [Tilletiaria anomala UBC
951]
MPETREDSVYLAKLAEQAERYEEMVENMKRVASSDQELTVEERNLLSVAYKNVIGARRASWRIVSSIEQKEESKGNE
AQVAMIKKYREKIEAELAKICEDILDVLDKHLIPSAASGESKVFYHKMKGDYHRYLAEFATGDKRKDSADKSLEAYK
AASDVAVTELPPTHPIRLGLALNFSVFYYEILNSPDRACHLAKQAFDDAIAELDTLSEESYKDSTLIMQLLRDNLTL
WTSDMQDSEAKAEPAAGEQPAPAEAAKGEEAA
>gi|761759007|emb|CEP63288.1| LALA0S07e06722g1_1 [Lachancea lanzarotensis]
MSQSREDSVYLAKLAEQAERYEEMVDSMKAVASSGQELSVEERNLLSVAYKNVIGARRASWRIVSSIEQKEEAKEKS
EHQVKLIRNYRSKIESELTKICDDILSVLDTHLIPSATTGESKVFYYKMKGDYHRYLAEFSSGEVRDGATNASLEAY
KTASEIATTELPPTHPIRLGLALNFSVFYYEIQNSPDKACHLAKQAFDDAIAELDTLSEESYKDSTLIMQLLRDNLT
LWTSDMSEAGQDEQQPAEGAQE
>gi|255718733|ref|XP_002555647.1| KLTH0G14146p [Lachancea thermotolerans]
MSQSREDSVYLAKLAEQAERYEEMVDSMKAVASSGQELSVEERNLLSVAYKNVIGARRASWRIVSSIEQKEEAKDKS
EHQVKLIRDYRSKIETELTKICDDILSVLDTHLIPSATTGESKVFYYKMKGDYHRYLAEFSSGEVRDKATNASLEAY
KTASEIATTELPPTHPIRLGLALNFSVFYYEIQNSPDKACHLAKQAFDDAIAELDTLSEESYKDSTLIMQLLRDNLT
LWTSDMSEAGQDEQQPAEGAQE
>gi|61676645|gb|AAX51846.1| 14-3-3 protein [Paxillus involutus]
MPESREDSVYLAKLAEQAERYEEMVENMKRVASSDQELTVEERNLLSVAYKNVIGARRASWRIVSSIEQKEESKGNE
AQVAMIKGYREKIETELAKICEDILDVLDKHLIPSAASGESKVFYHKMMGDYHRYLAEFATGDKRKDSADKSLAAYK
DASDVAVTELPPTHPIRLGLALNFSVFYYEILNSPDRACHLAKQAFDDAIAELDTLSEESYKDSTLIMQLLRDNLTL
WTSDMQDTDKTDSKDEGENAGDD
>gi|384490975|gb|EIE82171.1| 14-3-3 family protein epsilon [Rhizopus delemar
RA 99-880]
MATERTDKVYMAKIAEQAERYDEMVSYMKEVAKMGVDLTVEERNLLSVAYKNVIGARRASWRIVSSIEQKEESKGHE
TQVTKIKEYRNKIENELYEVCADILALLSDHLIPAAGDGEAKVFYYKMKGDYHRYVAEYATGDERKDAANLAHEAYK
KATEVAQVELATTHPIRLGLALNFSVFYYEILNSPDRACHLAKQAFDDAIAELDTLSEESYKDSTLIMQLLRDNLTL
WTSDLQEAESNDKPEEPKGPTDE
>gi|661182596|emb|CDH54915.1| 14-3-3 protein [Lichtheimia corymbifera
JMRC:FSU:9682]
MASERENDVYMAKLAEQAERYDEMVQYTKDVAKMGVELTVEERNLLSVAYKNVIGARRASWRIVSSIEQKEESKGNN
AQVEKIKVYRQKIENELKDVCSDILSVLKDNLIPNAQGGESKVFYYKMMGDYHRYLAEFLTGEARKESATQAHEAYK
TATDVAQTELAPTHPIRLGLALNFSVFYYEILNSPDRACHLAKQAFDDAIAELDTLSEESYKDSTLIMQLLRDNLTL
WTSDLQDTDNKGEQNEGEATEESK
>gi|302693879|ref|XP_003036618.1| 14-3-3 protein [Schizophyllum commune H4-8]
MPESREDSVYLAKLAEQAERYEEMVENMKRVASSDQELTVEERNLLSVAYKNVIGARRASWRIVSSIEQKEESKGNE
AQVSMIKGYREKIEGELAKICEDILDVLEKHLIPSAASGESKVFYHKMMGDYHRYLAEFATGDKRKDSADKSLEAYK
AASDVAVTELPPTHPIRLGLALNFSVFYYEILNSPDRACHLAKQAFDDAIAELDTLSEESYKDSTLIMQLLRDNLTL
WTSDMQDSGEKPADSKDEAPEAPGDE
>gi|831331195|gb|KLT40777.1| 14-3-3 protein [Trichosporon oleaginosus]
MSTSREDSVYLAKLAEQAERYEEMVENMKAVASSDKELTVEERNLLSVAYKNVIGARRASWRIVSSIEQKEESKGNE
AQVAMIKTYREKIEAELAKICKDILEVLDKHLIPSAASGESKVFYHKMMGDYHRYLAEFATGDKRKDSADKSLEAYK
AASDVAVTELPPTHPIRLGLALNFSVFYYEILNSPDRACHLAKQAFDDAIAELDTLSEESYKDSTLIMQLLRDNLTL
WTSDMQDSAEEPKKEEESKEEAA
>gi|384497424|gb|EIE87915.1| protein BMH2 [Rhizopus delemar RA 99-880]
MSTERENNVYMAKLAEQAERYDEMVTFTKDVAKMGVELTVEERNLLSVAYKNVIGARRASWRIVSSIEQKEESKGNS
TQVEKIKAYRQKIETELQDVCKDILAVLSENLIPNAQGGESKVFYYKMMGDYHRYLAEFLTSEARKESGNQAHEAYK
TATDIAQTELAPTHPIRLGLALNFSVFYYEISNSPDRACHLAKQAFDDAIAELDTLSEESYKDSTLIMQLLRDNLTL
WTSDLQEEADKVDDNKADATDDSK

>gi|599104601|ref|XP_007382290.1| 14-3-3 1 protein [Punctularia strigosozonata HHB-11173 SS5]
MTQSREDSVYLAKLAEQAERYEEMVENMKRVASSDQELTVEERNLLSVAYKNVIGARRASWRIVSSIEQKEESKGNE
AQVQMIKGYREKIESELAKICEDILAVLDKHLIPSAATGESKVFYHKMMGDYHRYLAEFATGDKRKESADKSLEAYK
AASDVAVTELPPTHPIRLGLALNFSVFYYEILNSPDRACHLAKQAFDDAIAELDTLSEESYKDSTLIMQLLRDNLTL
WTSDMQESDKPVDKEDNAEPADES
>gi|701778312|gb|KGQ11884.1| 14-3-3 protein [Beauveria bassiana D1-5]
MGNDDAVYLAKLAEQAERYEEMVENMKIVASEDRDLTVEERNLLSVAYKNVIGARRASWRIVTSIEQKEESKGNSSQ
VTLIKEYRQKIEDELAKICEDILDVLDKHLIPSAKSGESKVFYHKMKGDYHRYLAEFATGDKRKDSADKSLEAYKGA
TDVAQSELPPTHPIRLGLALNFSVFYYEILNAPDQACHLAKQAFDDAIAELDTLSEESYKDSTLIMQLLRDNLTLWT
SSEAETAPAAEGASKEEPVAAEEPKAEEPKAEEPKAAE
>gi|588260051|ref|XP_006959179.1| 14-3-3 protein [Wallemia mellicola CBS 633.66]
MTETREDHVYLAKLAEQAERYEEMVESMKKVASSDQELTVEERNLLSVAYKNVIGARRASWRIVSSIEQKEESKGND
SQVGMIKSYREKIELELAKICEDILAVLDGHLIPSAASGESKVFYHKMKGDYHRYLAEFATGDRRKGSAEASLEAYK
SASDVAITELPPTHPIRLGLALNFSVFYYEILNSPDRACHLAKQAFDDAIAELDTLSEESYKDSTLIMQLLRDNLTL
WTSDMQESDSKPEEKSDAKPEESAQPAQGEQA
>gi|661184855|emb|CDH53016.1| 14-3-3 family protein [Lichtheimia corymbifera JMRC:FSU:9682]
MSTEREDKVYMAKIAEQAERYDEMVTYMKEVAKQMSPDLSVEERNLLSVAYKNVIGARRASWRIVSSIEQKEESKGH
DAQVNKIKEYRKKIESELYDVCNDILELLKDCLIPAASDGEAKVFYYKMKGDYHRYVAEYATSEDRKTAASEAHEAY
KSATDIAQVELATTHPIRLGLALNFSVFYYEILNSPDRACHLAKQAFDDAIAELDTLSEESYKDSTLIMQLLRDNLT
LWTSDLQEDGDKQDKSTEAPKDNNEQAQE
>gi|170091548|ref|XP_001876996.1| 14-3-3 protein [Laccaria bicolor S238N-H82]
MPESREDSVYLAKLAEQAERYEEMVENMKRVASSDQELTVEERNLLSVAYKNVIGARRASWRIVSSIEQKEESKGNE
AQVSMIKGYREKIESELAKICEDILDVLDKHLIPSAASGESKVFYHKMMGDYHRYLAEFATGDKRKDSADKSLEAYK
AASDVAVTELPPTHPIRLGLALNFSVFYYEILNSPDRACHLAKQAFDDAIAELDTLSEESYKDSTLIMQLLRDNLTL
WTSDMQDSADKPSDGKDEAAEAPADD
>gi|630207454|ref|XP_007853881.1| 14-3-3 protein [Moniliophthora roreri MCA 2997]
MPESREDSVYLAKLAEQAERYEEMVENMKRVASSDQELTVEERNLLSVAYKNVIGARRASWRIVSSIEQKEESKGNE
AQVSMIKGYREKIESELAKICEDILDVLDKHLIPSAASGESKVFYHKMMGDYHRYLAEFATGDKRKDSADKSLEAYK
AASDVAVTELPPTHPIRLGLALNFSVFYYEILNSPDRACHLAKQAFDDAIAELDTLSEESYKDSTLIMQLLRDNLTL
WTSDMQDSDKPADKEDAGDAPADD
>gi|568442830|ref|XP_006456745.1| 14-3-3 protein [Agaricus bisporus var. bisporus H97]
MQMTTEVREDAVYLAKLAEQAERYEEMVENMKRVASSDQELTVEERNLLSVAYKNVIGARRASWRIVSSIEQKEETK
GNETQVRMIKSYREKIEGELADICEDILDVLDKHLIPSAATGESKVFYHKMMGDYHRYLAEFATGDKRKDSADKSLE
AYKAASDVAVTELPPTHPIRLGLALNFSVFYYEILNSPDRACHLAKQAFDDAIAELDTLSEESYKDSTLIMQLLRDN
LTLWTSDMQEAESATAENKETEATDAQPTDD
>gi|597987253|ref|XP_007365004.1| 14-3-3-domain-containing protein [Dichomitus squalens LYAD-421 SS1]
MAAHANNPLPSSEFNFFIYLPSRLASTLSSSARHSYRHPFTLPPPHTNTIAMAQSREDSVYLAKLAEQAERYEEMVE
NMKRVASSDQELTVEERNLLSVAYKNVIGARRASWRIVSSIEQKEESKGNEVQVKMIKGYREKIESELAKICEDILD
VLDKHLIPSAASGESKVFYHKMMGDYHRYLAEFATGDKRKESADKSLEAYKAASDVAVTELPPTHPIRLGLALNFSV
FYYEILNSPDRACHLAKQAFDDAIAELDTLSEESYKDSTLIMQLLRDNLTLWTSDMQESDKPVDKDDAADAPAEEGA
>gi|19115079|ref|NP_594167.1| 14-3-3 protein Rad24 [Schizosaccharomyces pombe 972h-]
MSTTSREDAVYLAKLAEQAERYEGMVENMKSVASTDQELTVEERNLLSVAYKNVIGARRASWRIVSSIEQKEESKGN
TAQVELIKEYRQKIEQELDTICQDILTVLEKHLIPNAASAESKVFYYKMKGDYYRYLAEFAVGEKRQHSADQSLEGY
KAASEIATAELAPTHPIRLGLALNFSVFYYEILNSPDRACYLAKQAFDEAISELDSLSEESYKDSTLIMQLLRDNLT
LWTSDAEYSAAAAGGNTEGAQENAPSNAPEGEAEPKADA
>gi|1040756|emb|CAA55795.1| rad24 [Schizosaccharomyces pombe]
MSTTSREDAVYLAKLAEQAERYEGMVENMKSVASTDQELTVEERNLLSVAYKNVIGARRASWRIVSSIEQKEESKGN
TAQVELIKEYRQKIEQELDTICQDILTVLEKHLIPNAASAESKVFYYKMKGDYYRYLAEFAVGEKRQHSADQSLEGY
KAASEIATAELAPTHPIRLGLALNFSVFYYEILNSPDRACYLAKQAFDEAISELDSLSEESYKDSTLIMQLLRDNLT
LWTSDAEYSAAAAGGNTEGAQENAPSNAPEGEREPKATHR

>gi|751837405|emb|CEL63931.1| 14-3-3 protein homolog OS=Trichoderma harzianum PE=2 SV=1 [Rhizoctonia solani AG-1 IB]
MWVEGAGWGSGLNRGRCDLDRPLPSESVVSALVNCGVYNPSLSLLYSLYIIMSSSREDSVYLAKLAEQAERYEEMVE
NMKRVASSDQELTVEERNLLSVAYKNVIGARRASWRIVSSIEQKEESKGNDAQVTMIKGYREKIEAELAKICEDILD
VLDKHLIPSAASGESKVFYHKMMGDYHRYLAEFATGDKRKESADKSLEAYKSASDVAITELPPTHPIRLGLALNFSV
FYYEILNSPDRACHLAKQAFDDAIAELDTLSEESYKDSTLIMQLLRDNLTLWTSDMQDSADKSGDKDEAPAEADDAS
KA
>gi|660963601|gb|KEP48927.1| DNA damage checkpoint protein rad24 [Rhizoctonia solani 123E]
MTDSREDSVYLAKLAEQAERYEEMVENMKRVASSDQELTVEERNLLSVAYKNVIGARRASWRIVSSIEQKEESKGND
AQVTMIKGYREKIETELAKICEDILDVLDKHLIPSAASGESKVFYHKMMGDYHRYLAEFATGDKRKQSADKSLDAYK
AASDVAVTELPPTHPIRLGLALNFSVFYYEILNSPDRACHLAKQAFDDAIAELDTLSEESYKDSTLIMQLLRDNLTL
WTSDMQDSADKSGDKDDAPAEAGDADKA
>gi|661184856|emb|CDH53017.1| 14-3-3 family protein [Lichtheimia corymbifera JMRC:FSU:9682]
MSTEREDKVYMAKIAEQAERYDEMVTYMKEVAKMSPDLSVEERNLLSVAYKNVIGARRASWRIVSSIEQKEESKGHD
AQVNKIKEYRKKIESELYDVCNDILELLKDCLIPAASDGEAKVFYYKMKGDYHRYVAEYATSEDRKTAASEAHEAYK
SATDIAQVELATTHPIRLGLALNFSVFYYEILNSPDRACHLAKQAFDDAIAELDTLSEESYKDSTLIMQLLRDNLTL
WTSDLQEDGDKQDKSTEAPKDNNEQAQE

#### 3. Homology of *Plasmodium falciparum 3D7* 14-3-3I protein with 14-3-3 proteins from Animals

**Query Sequence:** 14-3-3I protein, putative, *Plasmodium falciparum 3D7*; MAL8P1.69
**Subject:** Animals (taxid:33208)

##### Protein sequences showing homology with 14-3-3I, *Plasmodium falciparum 3D7*

>gi|148298752|ref|NP_001091764.1| 14-3-3 epsilon protein [Bombyx mori]
MSEREDNVYKAKLAEQAERYDEMVEAMKNVASRNVSDNELTVEERNLLSVAYKNVIGARRASWRIISSIEQKEETKG
AEDKLNMIRAYRSQVEKELRDICSDILGVLDKYLIPSSQTGESKVFYYKMKGDYHRYLAEFATGNDRKEAAENSLVA
YKAASDIAMTELPPTHPIRLGLALNFSVFYYEILNSPDRACRLAKAAFDDAIAELDTLSEESYKDSTLIMQLLRDNL
TLWTSDMQGDGESADAEQKEPAQDGEDQDVS
>gi|357618137|gb|EHJ71231.1| 14-3-3 epsilon protein [Danaus plexippus]
MSEREDNVYKAKLAEQAERYDEMVEAMKNVASRNISDNELTVEERNLLSVAYKNVIGARRASWRIISSIEQKEETKG
AEDKLSMIRAYRSQVEKELRDICSDILGVLDKHLIPASQTGESKVFYYKMKGDYHRYLAEFATGNDRKEAAENSLVA
YKAASDIAMTELPPTHPIRLGLALNFSVFYYEILNSPDRACRLAKAAFDDAIAELDTLSEESYKDSTLIMQLLRDNL
TLWTSDMQGDAEPDQEPKEPAQEPDDQDVS
>gi|822092769|ref|NP_001296019.1| 14-3-3 protein epsilon [Plutella xylostella]
MSEREDNVYKAKLAEQAERYDEMVEAMKNVASRNVSDNELTVEERNLLSVAYKNVIGARRASWRIISSIEQKEETKG
AEDKLSMIRAYRSQVEKELRDICADILAVLDKHLIPSSQTGESKVFYYKMKGDYHRYLAEFATGNDRKEAAEHSLVA
YKAASEIAMTELPPTHPIRLGLALNFSVFYYEILNSPDRACRLAKAAFDDAIAELDTLSEESYKDSTLIMQLLRDNL
TLWTSDMQGDGESGEAEQKEQAQDVEDQDVS
>gi|765527265|gb|AJS10721.1| 14-3-3 epsilon [Tenebrio molitor]
MSEREDNVYKAKLAEQAERYDEMVDAMKKVAKLDLELTVEERNLLSVAYKNVIGARRASWRIISSIEQKEESKGTDD
KLEMIRQYRSQVEKELRDICSDILTVLDKHLIPAASTGESKVFYYKMKGDYHRYLAEFATGNDRKDAAEHSLVAYKS
ASDIAMTELPPTHPIRLGLALNFSVFYYEILNSPDRACRLAKAAFDDAIAELDTLSEESYKDSTLIMQLLRDNLTLW
TSDMQGDGEAEPKEQLQDVEDQDVS
>gi|359843276|gb|AEV89773.1| 14-3-3 protein epsilon [Schistocerca gregaria]
MSERDDNVYKAKLAEQAERYDEMVEAMKKVASLDVELTVEERNLLSVAYKNVIGARRASWRIISSIEQKEENKGAEE
KLEMIRGYRSQVEKELKDICSDILGVLDKHLIPCASTGESKVFYYKMKGDYHRYLAEFATGNDRKEAAENSLVAYKA
ASDIAMTELPPTHPIRLGLALNFSVFYYEILNSPERACRLAKAAFDDAIAELDTLSEESYKDSTLIMQLLRDNLTLW
TSDMQGDGETEQKEQLQDVEDQDVS

>gi|56118781|ref|NP_001008156.1| 14-3-3 protein epsilon [Xenopus (Silurana) tropicalis]
MEEREDLVYRAKLAEQAERYDEMVESMKKVAGMDVELTVEERNLLSVAYKNVIGARRASWRIISSIEQKEENKGGED
KLKMIREYRQMVEAELKSICNDILDVLDKHLIPAANSGESKVFYYKMKGDYHRYLAEFAQGNDRKEAAENSLVAYKA
ASDIAMTELPPTHPIRLGLALNFSVFYYEILNSPDRACRLAKAAFDDAIAELDTLSEESYKDSTLIMQLLRDNLTLW
TSDMQGDGEEQNKDALQDVEDENQ
>gi|675377550|gb|KFM70452.1| 14-3-3 protein epsilon [Stegodyphus mimosarum]
MADREDNVYKAKLAEQAERYDEMVEAMKKVASLDLELTVEERNLLSVAYKNVIGARRASWRIISSIEQKEENKGAEN
RLEMIKTYRVQVETELKDICQDILDVLDKHLIPTASTGESKVFYYKMKGDYHRYLAEFATGNDRKEAAENSLVAYKA
ASDIAMTELPPTHPIRLGLALNFSVFYYEILNSPDRACRLAKAAFDDAIAELDTLSEESYKDSTLIMQLLRDNLTLW
TSDMQHDGESEQKEQVQDVEDQDVS
>gi|927544317|gb|ALE20564.1| 14-3-3 protein epsilon [Leptinotarsa decemlineata]
MSEREDNVYKAKLAEQAERYDEMVEAMKKVAKLDLELTVEERNLLSVAYKNVIGARRASWRIISSIEQKEESKGTDD
KLEMIRQYRSQVEKELRDICSDILSVLDKHLIPAASSGESKVFYYKMKGDYHRYLAEFATGNDRKDAAEHSLVAYKS
ASDIAMTELPPTHPIRLGLALNFSVFYYEILNSPDRACRLAKAAFDDAIAELDTLSEESYKDSTLIMQLLRDNLTLW
TSDMQGDGEAEPKEQLQDVEDQDVS
>gi|212283378|gb|ACJ23184.1| 14-3-3 epsilon [Penaeus monodon]
MTDREDNVYRAKLAEQAERYDEMVDAMKLVASMDVELTVEERNLLSVAYKNVIGARRASWRIISSIEQKEENKGGEE
KLEMIRNYRTQVEKELKDICSDILGLLDKHLIPTASAGESRVFFYKMKGDYHRYLAEFATGNDRKAAAENSLVAYKA
ASDIAMTELPPTHPIRLGLALNFSVFYYEILNSPERACRLAKAAFDDAIAELDTLSEESYKDSTLIMQLLRDNLTLW
TSDMQGEASKGTVPDASLCEAARRLCRDQICISPNIPESTLV
>gi|148225538|ref|NP_001080705.1| tyrosine 3-monooxygenase/tryptophan 5-monooxygenase activation protein, epsilon [Xenopus laevis]
MEEREDLVYRAKLAEQAERYDEMVESMKKVAGMDVELTVEERNLLSVAYKNVIGARRASWRIISSIEQKEENKGGED
KLKMIREYRQMVETELKSICNDILDVLDKHLIPAASSGESKVFYYKMKGDYHRYLAEFAQGNDRKEAAENSLVAYKA
ASDIAMTELPPTHPIRLGLALNFSVFYYEILNSPDRACRLAKAAFDDAIAELDTLSEESYKDSTLIMQLLRDNLTLW
TSDMQGDGEDQNKEALQDVEDENQ
>gi|61651838|ref|NP_001013359.1| tyrosine 3-monooxygenase/tryptophan 5-monooxygenase activation protein, epsilon polypeptide 2 [Danio rerio]
MADREHLVYQAKLAEQAERYDEMVESMKNVAGKDEDLSVEERNLLSVAYKNVIGARRASWRIISSIEQKEESKGGAD
KLKMIREYRQTVENELKSICNDILDVLDKHLIPAANTGESKVFYYKMKGDYHRYLAEFATGNDRKEAAENSLVAYKA
ASDIAMTELPPTHPIRLGLALNFSVFYYEILNSPDRACRLAKAAFDDAIAELDTLSEDSYKDSTLIMQLLRDNLTLW
TSDIQGDGEEQSKTAPQDAEEEKQ
>gi|308321454|gb|ADO27878.1| 14-3-3 protein epsilon [Ictalurus furcatus]
MDRDHFVYQAKLAEQAERYDEMVESMKSVAGKDVELTVEERNLLSVAYKNVIGARRASWRIISSIEQKEESKGGEDK
LKMIREYRQTVEKELKSICNDILDVLDKHLIPSANTGESKVFYYKMKGDYHRYLAEFATGNDRKEAAENSLVAYKAA
SDIAMIELPPTHPIRLGLALNFSVFYYEILNSPDRACRLAKAAFDDAIAELDTLSEESYKDSTLIMQLLRDNLTLWT
SDMQGDGEEQDKEALQDVEDENQ
>gi|185134340|ref|NP_001117944.1| 14-3-3E1 protein [Oncorhynchus mykiss]
MGDRDELVYQAKLAEQAERYDEMVESMKRVAGLDVELTVEERNLLSVAYKNVIGARRASWRIISSIEQREENKGGED
KLKMIREYRQTVENELKSICNDILDVLDKHLIPAANTGESKVFYYKMKGDYHRYLAEFATGNDRKEAAENSLVAYKA
ASDIAMIELPPTHPIRLGLALNFSVFYYEILNSPDRACRLAKAAFDDAIAELDTLSEESYKDSTLIMQLLRDNLTLW
TSDMQGDAGEEQQNKEALQDVEDEPQ
>gi|501293543|dbj|BAN20616.1| 14-3-3 epsilon protein [Riptortus pedestris]
MSEREDNVYKAKLAEQAERYDEMVEAMKLVASLDLELTVEERNLLSVAYKNVIGARRASWRIISSIEQKEENKGADD
KLDMIRQYRSQVEKELREICSDILGVLDKHLIPCASTGESKVFYYKMKGDYHRYLAEFATGNDRKEAAENSLVAYKA
ASDIAMTELPPTHPIRLGLALNFSVFYYEILNSPDRACRLAKAAFDDAIAELDTLSEESYKDSTLIMQLLRDNLTLW
TSDMQGDGNSEQKAELQDVEDQDVS
>gi|194764555|ref|XP_001964394.1| GF23073 [Drosophila ananassae]
MTERENNVYKAKLAEQAERYDEMVEAMKKVASMDVELTVEERNLLSVAYKNVIGARRASWRIITSIEQKEETKGAEE
KLEMIKTYRGQVEKELRDICSDILNVLEKHLIPCATTGESKVFYYKMKGDYHRYLAEFATGSDRKDAAEKSLIAYKE
ASDIAMNDLPPTHPIRLGLALNFSVFYYEILNSPDRACRLAKAAFDDAIAELDTLSEESYKDSTLIMQLLRDNLTLW
TSDMQAEEVDPNTGDGEPKDTQIADPEEQDVS
>gi|924563216|gb|ALC47874.1| 14-3-3epsilon [Drosophila busckii]
MTERENNVYKAKLAEQAERYDEMVEAMKKVASMDVELTVEERNLLSVAYKNVIGARRASWRIITSIEQKEENKGAEE
KLDMIKTYRGQVEKELRDICSDILNVLEKHLIPCATSGESKVFYYKMKGDYHRYLAEFATGSDRKDAAENSLIAYKA

ASDIAMNDLPPTHPIRLGLALNFSVFYYEILNSPDRACRLAKAAFDDAIAELDTLSEESYKDSTLIMQLLRDNLTLW
TSDMQADDSTTGDGETKQEIQDVEDQDVS
>gi|195036824|ref|XP_001989868.1| GH18560 [Drosophila grimshawi]
MTERENNVYKAKLAEQAERYDEMVEAMKKVASMDVELTVEERNLLSVAYKNVIGARRASWRIITSIEQKEENKGAEE
KLEMIKTYRGQVEKELRDICSDILNVLEKHLIPCATSGESKVFYYKMKGDYHRYLAEFATGSDRKDAAENSLIAYKA
ASDIAMNDLPPTHPIRLGLALNFSVFYYEILNSPDRACRLAKAAFDDAIAELDTLSEESYKDSTLIMQLLRDNLTLW
TSDMQADDSTTGDGEPKQEIQDVEDQDVS
>gi|290562958|gb|ADD38873.1| 14-3-3 protein epsilon [Lepeophtheirus salmonis]
MAEREDCVYKAKLAEQAERYDEMVTSMKMVASMDLELTVEERNLLSVAYKNVIGARRASWRIISSLEAKEGNKASED
KLNLIKNYRTQVEKELKDICGDILSVLDKHLIPCSNTGESKVFYYKMKGDYHRYLAEFATNNDRKEAAENSLVAYKA
ASDTAMSELPTTHPIRLGLALNFSVFYYEILNSPDRACRLAKAAFDDAIAELDTLSEESYKDSTLIMQLLRDNLTLW
TSDMQAEDGDGEKKEKVEDVESGNAEAQPEAAAS
>gi|24647885|ref|NP_732309.1| 14-3-3epsilon, isoform A [Drosophila
melanogaster]
MTERENNVYKAKLAEQAERYDEMVEAMKKVASMDVELTVEERNLLSVAYKNVIGARRASWRIITSIEQKEENKGAEE
KLEMIKTYRGQVEKELRDICSDILNVLEKHLIPCATSGESKVFYYKMKGDYHRYLAEFATGSDRKDAAENSLIAYKA
ASDIAMNDLPPTHPIRLGLALNFSVFYYEILNSPDRACRLAKAAFDDAIAELDTLSEESYKDSTLIMQLLRDNLTLW
TSDMQAEEVDPNAGDGEPKEQIQDVEDQDVS
>gi|240849453|ref|NP_001155476.1| 14-3-3 protein epsilon [Acyrthosiphon
pisum]
MSEREENVYKAKLAEQAERYDEMVESMKKVASLDVELSVEERNLLSVAYKNVIGARRASWRIISSIEQKEENKGAEE
KLEMIRQYRSQVEKELRDICSDILNVLDKHLIACAASGESKVFYYKMKGDYHRYLAEFATGDDRKEAAEHSLVAYKA
ASDIANQDLPPTHPIRLGLALNFSVFYYEILNTPDRACHLAKKAFDEAIAELDTLSEESYKDSTLIMQLLRDNLTLW
TSDMQGDGTEAEPKEQLQDVEDQDVS
>gi|47086819|ref|NP_997770.1| 14-3-3 protein epsilon [Danio rerio]
MGDREDLVYQAKLAEQAERYDEMVDSMKKVAGMDVELTVEERNLLSVAYKNVIGARRASWRIISSIEQKEENKGGED
KLKMIREYRQTVENELKSICNDILDVLDKHLIPAANSGESKVFYYKMKGDYHRYLAEFATGNDRKEAAENSLVAYKA
ASDIAMTDLQPTHPIRLGLALNFSVFYYEILNSPDRACRLAKAAFDDAIAELDTLSEESYKDSTLIMQLLRDNLTLW
TSDMQGDGEEQNKEALQDVEDENQ
>gi|5803225|ref|NP_006752.1| 14-3-3 protein epsilon [Homo sapiens]
MDDREDLVYQAKLAEQAERYDEMVESMKKVAGMDVELTVEERNLLSVAYKNVIGARRASWRIISSIEQKEENKGGED
KLKMIREYRQMVETELKLICCDILDVLDKHLIPAANTGESKVFYYKMKGDYHRYLAEFATGNDRKEAAENSLVAYKA
ASDIAMTELPPTHPIRLGLALNFSVFYYEILNSPDRACRLAKAAFDDAIAELDTLSEESYKDSTLIMQLLRDNLTLW
TSDMQGDGEEQNKEALQDVEDENQ
>gi|225712634|gb|ACO12163.1| 14-3-3 protein epsilon [Lepeophtheirus salmonis]
MAEREDCVYKAKLAEQAERYDEMVTSMKMVASMDLELTVEERNLLSVAYKNVIGARRASWRIISSLEAKEGNKASED
KLNLIKNYRTQVEKELKDICGDILSVLDKHLIPCSNTGESKVFYYKMKGDYHRYLAEFATNNDRKEAAENSLVAYKA
ASDTAMSELPTTHPIRLGLALNFSVFYYEILNSPDRACRLAKAAFDDVIAELDTLSEESYKDSTLIMQLLRDNLTLW
TSDMQAEDGDGEKKEKVEDVESGNAEAQPEAAAS
>gi|24647891|ref|NP_732312.1| 14-3-3epsilon, isoform C [Drosophila
melanogaster]
MTERENNVYKAKLAEQAERYDEMVEAMKKVASMDVELTVEERNLLSVAYKNVIGARRASWRIITSIEQKEENKGAEE
KLEMIKTYRGQVEKELRDICSDILNVLEKHLIPCATSGESKVFYYKMKGDYHRYLAEFATGSDRKDAAENSLIAYKA
ASDIAMNDLPPTHPIRLGLALNFSVFYYEILNSPDRACRLAKAAFDDAIAELDTLSEESYKDSTLIMQLLRDNLTLW
TSDMQAEGDGEPKEQIQDVEDQDVS
>gi|288784881|gb|ADC53751.1| AT09839p [Drosophila melanogaster]
FNTMTERENNVYKAKLAEQAERYDEMVEAMKKVASMDVELTVEERNLLSVAYKNVIGARRASWRIITSIEQKEENKG
AEEKLEMIKTYRGQVEKELRDICSDILNVLEKHLIPCATSGESKVFYYKMKGDYHRYLAEFATGSDRKDAAENSLIA
YKAASDIAMNDLPPTHPIRLGLALNFSVFYYEILNSPDRACRLAKAAFDDAIAELDTLSEESYKDSTLIMQLLRDNL
TLWTSDMQAEGDGEPKEQIQDVEDQDVS
>gi|24647889|ref|NP_732311.1| 14-3-3epsilon, isoform D [Drosophila
melanogaster]
MTERENNVYKAKLAEQAERYDEMVEAMKKVASMDVELTVEERNLLSVAYKNVIGARRASWRIITSIEQKEENKGAEE
KLEMIKTYRGQVEKELRDICSDILNVLEKHLIPCATSGESKVFYYKMKGDYHRYLAEFATGSDRKDAAENSLIAYKA
ASDIAMNDLPPTHPIRLGLALNFSVFYYEILNSPDRACRLAKAAFDDAIAELDTLSEESYKDSTLIMQLLRDNLTLW
TSDMQAEDPNAGDGEPKEQIQDVEDQDVS
>gi|925671797|gb|KOX69543.1| 14-3-3 protein epsilon [Melipona quadrifasciata]

MSEREDNVYKAKLAEQAERYDEMVEAMKKVASLDVELTVEERNLLSVAYKNVIGARRASWRIISSIEQKEENKGAER
KLEMIRQYRSQVEKELKDICADILGVLDKHLLPCASTGESKVFYYKMKGDYHRYLAEFAVGNDRKEAAENSLVAYKA
ASDTAMTDLPPTHPIRLGLALNFSVFYYEILNSPDRACRLAKAAFDDAIAELDTLSEESYKDSTLIMQLLRDNLTLW
TSDMQGDG
>gi|24647887|ref|NP_732310.1| 14-3-3epsilon, isoform B [Drosophila
melanogaster]
MTERENNVYKAKLAEQAERYDEMVEAMKKVASMDVELTVEERNLLSVAYKNVIGARRASWRIITSIEQKEENKGAEE
KLEMIKTYRGQVEKELRDICSDILNVLEKHLIPCATSGESKVFYYKMKGDYHRYLAEFATGSDRKDAAENSLIAYKA
ASDIAMNDLPPTHPIRLGLALNFSVFYYEILNSPDRACRLAKAAFDDAIAELDTLSEESYKDSTLIMQLLRDNLTLW
TSDMQAEVDPNAGDGEPKEQIQDVEDQDVS
>gi|195450008|ref|XP_002072323.1| GK22388 [Drosophila willistoni]
MTERENNVYKAKLAEQAERYDEMVEAMKKVASMDVELTVEERNLLSVAYKNVIGARRASWRIITSIEQKEENKGAEE
KLEMIKTYRGQVEKELRDICSDILNVLEKHLIPCATSGESKVFYYKMKGDYHRYLAEFATGSDRKDAAENSLIAYKA
ASDIAMNDLPPTHPIRLGLALNFSVFYYEILNSPDRACRLAKAAFDDAIAELDTLSEESYKDSTLIMQLLRDNLTLW
TSDMQAEDPNAGDGEPKGEIQDVEDQDVS
